# Assisted colonisation prospects for the black-veined white butterfly in England

**DOI:** 10.1101/2024.05.21.595182

**Authors:** Chris D Thomas, Charles A Cunningham, Neil A C Hulme, Eleanor C Corrigan, Bonnie Metherell, Penny Green, Matthew Oates

## Abstract

The black-veined white butterfly, *Aporia crataegi*, reached the north-western edge of its European geographic distribution in the British Isles in the 19^th^ century, but became extinct in the early 20^th^ century, following several cold decades. Substantial areas of potential breeding habitat in southern Britain are currently available to this species, which requires scattered hawthorn (*Crataegus monogyna*) and suckering blackthorn scrub (*Prunus spinosa*), including infrequently cut hedgerows. These habitats are needed at a relatively large scale as the butterfly occurs as networks of colonies (metapopulations), ranging over large tracts of connected landscape. A number of events have increased habitat availability over the past 70 years (myxomatosis reduced rabbit populations, which permitted host plant scrub regeneration; hedgerow management policies reduced cutting frequencies; rewilding and landscape connectivity initiatives are resulting in additional scrub). However, while *A. crataegi* males occasionally disperse several kilometres, it is unlikely that *A. crataegi* females will cross the English Channel in sufficient numbers to establish populations in southern England, without assistance.

Here, we (i) provide a review of the literature on species interactions, the habitat requirements, distribution and dispersal of *A. crataegi* and (ii) provide evidence that southern, eastern and central England are likely to be climatically suitable for reintroductions of *A. crataegi*. Substantial areas of England are already expected to be amongst the climatically most suitable parts of Europe for this insect. We identify a landscape of at least 100 km^2^ containing multiple patches of suitable habitats, and highlight co-benefits for other species that inhabit scrubland and successional mosaics. We use a climate-matching approach to assess climatically-similar locations to obtain source material most likely to establish in Britain. Areas of northern France and mid elevations in the Iberian Peninsula, including in the Pyrenees, provide potential suitable source locations due to close climatic matching and a large number of species records. We recommend reestablishment from more than one source, providing genetic diversity in the reintroduced population, enabling subsequent local adaptation to British conditions.

We highlight the opportunity for monitored releases to be undertaken within the landscape highlighted here, so as to evaluate population growth, host plant use, and rates of colonisation away from release sites that differ in their management, habitat, host plant characteristics, and proximity to other sites. This approach would aim to develop knowledge as a ‘living lab’ to inform future ‘best practice’ releases. In conclusion, the black-veined white, *A. crataegi*, has potential to become a model species for assisted colonisation projects where natural and human-created barriers have prevented range expansion into regions where the 21^st^ century climate is suitable for a species.

## INTRODUCTION

Restoring or reintroducing species that became extinct a substantial period of time ago and introducing or reintroducing species to regions in the context of climate change (assisted colonisation) will always involve a degree of uncertainty. The commonest questions are ‘*why did the species become extinct historically?*’, ‘*have those previous causes of decline been removed in the areas of proposed reintroduction?*’, and ‘*what might the impacts on other species be?*’, the latter being most important in the case of introductions to regions where a species did not occur historically. This approach is likely to become increasingly important as the distributions of species respond to climate change, but may not be able to achieve their full potential distributions because of dispersal barriers (Hoegh-Guldberg *et al*. 2008; Thomas 2011; IUCN/SSC 2013).

Here we consider options to reintroduce the black-veined white butterfly, *Aporia crataegi*, to Britain. This species was last recorded in southern England in the early to mid-1920s. Given the antiquity of the disappearance of this species, there is bound to be a degree of uncertainty about the exact combination of factors that led to its original decline. The main hypothesis is that the British climate was unsuitable for the species in the late 19^th^ and early 20^th^ centuries (Allan 1948; Pratt 1983; Eeles 2023). In the context of this historical uncertainty, the case for reintroduction to Britain requires consideration of:

i. whether the current and future climatic conditions are likely to be suitable for establishment,
ii. the likelihood that suitable habitats are available at sufficient scale, and
iii. what potential interactions with other species in Britain might take place during and subsequent to reestablishment.

Following a short background of the disappearance of *Aporia crataegi* from Britain, we consider these three issues in reverse order.

### A brief history

The last records of *Aporia crataegi* in the UK were from the early 1920s, both 1923 and 1925 being listed as possible years of the last sighting (Pratt 1983; Eeles 2023). However, there remains some debate whether *A. crataegi* still had a viable population in the preceding decades, the species having always been prone to extreme population fluctuations, including crashes in response to unsuitable climatic conditions (e.g., Ubach *et al*. 2022). Allan (1948) suggested that the persistence of this species in Britain beyond 1880 was due to its temporary reestablishment from imported continental stock, the butterfly’s decline over the preceding decades having been noted at the Entomological Society of London in 1884. Either way, populations declined and the distribution shrank severely in the mid to late 19^th^ century, regardless of whether full extinction of the ‘original’ British populations took place in the late 19^th^ century, or lingered on locally until the 1920s. Drawing on Pratt (1983) and Eeles (2023), and UK Biological Records Centre data, it appears that the butterfly ranged from South East England to the New Forest in southern-central England, and to the lower Severn valley region further to the west, but colonies were recorded as far north as Yorkshire (Figure 1; showing British records with sufficient spatial precision to plot). Extinction from the UK has primarily been attributed to a deterioration in climatic suitability, including a run of wet Septembers; however, specific weather events such as these would be unlikely to cause extinction unless the background climatic conditions were already marginal (see climatic analysis, below). While there are signs from museum specimens of genetic erosion in *A. crataegi* prior to its extinction in Britain (Whitla *et al*. 2023), this seems likely to be a *consequence* of the butterfly’s decline to low numbers, rather than the primary *cause* of decline. A number of small-scale reintroductions have been attempted but none was successful. A lack of any systematic monitoring or examination of these means that it is not possible to deduce why they failed.

**Figure 1.**
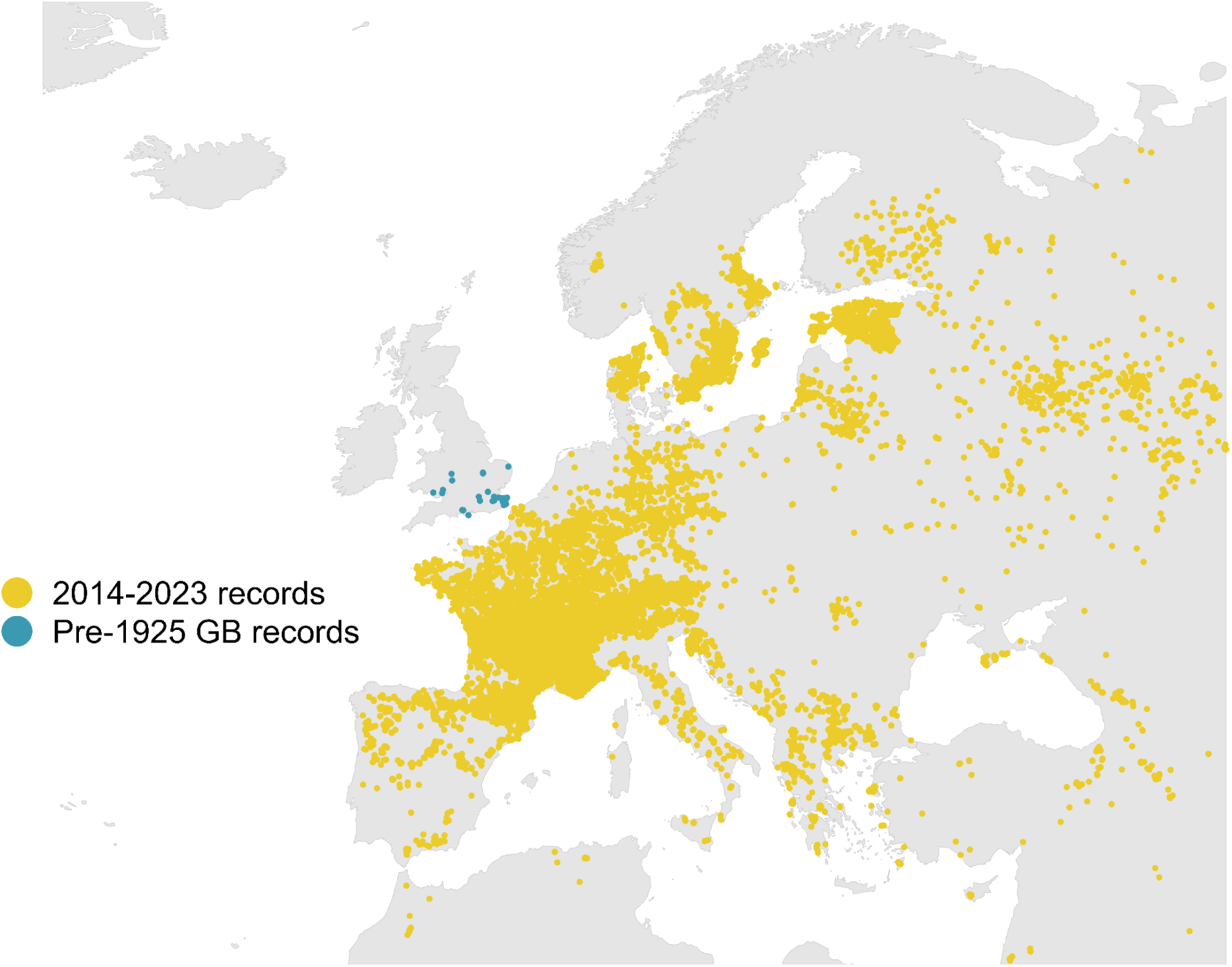
Black-veined white records from GBIF. All continental and African records from 2014-2023 are presented (yellow), with UK records prior to 1925 also shown (blue).

### New opportunities

With ongoing climate change, recent scientific studies suggest that much of southern Britain is now, and will remain, suitable for *A. crataegi* despite declines in many parts of Europe (Carroll *et al*., 2009). However, the species has declined in recent decades in the ‘near-continent’, primarily because of land use intensification. It is listed as extinct from the Netherlands and Czechia (Van Swaay *et al*., 2010), it no longer has breeding populations in Flanders (coastal regions of Belgium; Maes *et al*. 2016), and it is generally rare in the agriculturally intensive parts of north (eastern) France. As such, the nearest continental populations are generally small, with intensively-farmed landscapes constraining its potential northwards range extension. Combined with the geographic barrier of the English Channel, this means that the butterfly is considered unlikely to recolonise Britain naturally, at least within the foreseeable future. Occasional individuals - nearly all males - may cross the Channel as vagrants, but they are unlikely to establish viable populations (see dispersal section, below).

Given these barriers to natural colonisation, *A. crataegi* is unlikely to establish in Britain without assisted translocation, despite projected climate suitability. *Aporia crataegi* is classified as Regionally Extinct based on 2001 IUCN guidelines in the JNCC-sponsored Red listing of British butterflies (Fox *et al*. 2010), updated in 2022 (Fox *et al*. 2022). Assisted colonisation would, in this instance, aim to reverse a historical extinction, as well as to provide population sources in southern Britain for subsequent northwards range extension (both assisted and through natural dispersal). The main historical distribution of this species in Britain was southern and eastern England, including the proposed reestablishment county of Sussex. In addition to restoring an individual species that has been declining elsewhere (above), there are potential habitat management co-benefits for species that occupy similar habitats (see Discussion). *Aporia crataegi* can also enhance pollination (it is known to pollinate orchids; Lind *et al*. 2007), and adult black-veined white butterflies visit gardens and sheltered paths in search of nectar sources. Thus, once reestablished, the species could potentially be relatively easily encountered by the public, achieving wellbeing benefits - an insect version of the red kite!

In terms of habitat, *A. crataegi* has gregarious larvae that feed primarily on isolated bushes of blackthorn *Prunus spinosa* and hawthorn *Crataegus monogyna* in warm microclimates. As a relatively mobile species in which a few individuals disperse several kilometres in each generation, these habitats must be distributed across landscapes that enable dispersing females to find locations for oviposition, and provide males and females with opportunities to mate and obtain nectar. For this reason, we focus here on a substantial area of landscape in West Sussex, England, where it appears that sufficient landscape connectivity (of suitable habitats and microclimates) likely exists to support a viable reintroduced metapopulation, from which further British metapopulations may subsequently be established. As a butterfly species associated with scattered and ‘edge’ host plants that are shrubs or small trees, *A. crataegi* can potentially be adopted as an indicator of early-mid successional environments, on which many other threatened species also rely, as well as become an emblem for insect restoration and assisted colonisation.

Here, we consider the extent to which the reestablishment in Britain of *A. crataegi* from European populations (Figure 1) is a viable option.

## METHODS

This study combines literature review, field observation and assessment of the climate similarity of a key proposed reestablishment area in Britain to locations where *A. crataegi* populations currently exist in mainland Europe.

### Literature review

We searched Google Scholar for ‘*Aporia crataegi*’ (search date: 15 April 2024; beyond ‘hit 250’ no further relevant information was found) retaining all publications from Europe westwards of longitude 20°E (i.e., focussing on areas with a climatic match to southern Britain, Figure 9b). Potential racial/subspecies, biotic and climatic differences further eastwards make the relevance of such information questionable. Once publications were identified, citing literature was also scanned for relevance. Any further grey literature was sought from online searches, and considered when inspection of the information was considered to be ‘primary’ (as opposed to derived information in field guides and online accounts, which may include extensive lists of host plants from anywhere in the species’ Eurasian distribution, regardless of relevance to the region under consideration). The resulting literature was subdivided into publications indicating interactions with other species, those describing other aspects of the species habitat associations, and those evaluating the climatic requirements of the species.

### Field surveys

Selected sites were surveyed within an approximately 11 km x 9 km region in West Sussex, which represents a diversity of geologies, from the Wealden clays (at Knepp), through greensand (sandstone) to the chalk hills of the South Downs. We assessed the availability of scattered hawthorn (*Crataegus monogyna*) bushes and suckering blackthorn (*Prunus spinosa*) scrub or hedgerow edges, and the relative abundance of potential nectar plants with flowers in the blue/purple/red/pink spectrum. For the Knepp Estate, abundances of potential nectar sources were assessed using the DAFOR scale: D - Dominant >75% cover; A - Abundant 51-75%; F - Frequent 26-50%; O - Occasional 11-25%; R - Rare 1-10%. Overall DAFOR scores were calculated by summing across plant species. Based on prior expert knowledge of the landscape, the aim of this fieldwork was to identify the suitability of potential sites within the ∼11 km x 9 km landscape, rather than an assessment of the locations of *all* breeding habitats in the landscape.

#### Climatic suitability

We summarised the existing climatic suitability information from the literature, generated a new mean Spring and Summer measure of Central England temperatures since 1850 (National Climate Information Centre, 2024) and separately undertook a new climate similarity analysis of the match between the climate of the focal landscape and elsewhere in Europe.

For the climatic-match analysis, we used bioclimatic indicators for Europe downloaded from ECMWF Climate data store (available from <u>Copernicus</u>, Woulters 2021). These data are available at a 1 km x 1 km resolution as a mean for 1979-2018 for the region shown in Figure 1. Downloaded bioclimatic variables are:

- <u>Growing degree days</u> (K day year^-1^). The sum of daily degrees above the daily mean temperature of 278 K (5°C). The monthly data are aggregated over the months.
- <u>Annual precipitation</u> (mm year^-1^). This indicator corresponds to the official BIOCLIM variable BIO12, reflecting the annual mean of the daily precipitation *rate* (both liquid and solid phases). Given in units of m s^-1^, this was converted to total precipitation sum over the year, a conversion factor of 3600×24×365×1000, giving mm year^-1^ values.
- <u>Mean temperature of coldest month</u> (K). This was calculated by downloading monthly mean temperatures, and taking the minimum value.
- <u>Temperature seasonality</u> (K). Standard deviation of the monthly mean temperature multiplied by 100. This indicator corresponds to the official BIOCLIM variable BIO04.

To exclude marine areas from climate surfaces, national boundaries were downloaded using the *geodata* package (Hijmans *et al*. 2023) and climate variables were masked to terrestrial boundaries. The resulting surfaces are shown in Figure 7.

To identify climate analogues, we took six potential reintroduction sites in West Sussex (sites 1-6 in Tables 5, S1), close to the south coast of England, including the Knepp Estate and South Downs hills, extracted the climate values and took the mean. The absolute difference between the mean climate value of the reintroduction sites and all other climate values within the dataset was calculated for each climate variable. The most similar cells (20%, 30%, 40% and 50% quantiles) were identified and plotted for each bioclimatic variable (reintroduction sites as red circle; Figure 8). An overall climate similarity map was produced by identifying cells which were within specific quantiles for *all* of the climate variables (Figure 9a). Since moisture availability is unlikely to be a limiting factor for the host plants or for *A. crataegi* in Britain (whereas it is in e.g. the Mediterranean), we also estimated specific quantiles for *all* three of the temperature climate variables (Figure 9b).

To identify potential source populations, we downloaded *A. crataegi* presence records from GBIF (GBIF.org 2024). We carried out filtering of records by removing records that had; no coordinates, geospatial issues, zero counts, low coordinate precision (greater than or equal to 10km), or records with low verification confidence. We additionally cleaned the records using the *coordinate cleaner* package (Zizka *et al*. 2019), and removed UK records after 2000 as the species was confirmed extinct. All climatic suitability analysis was carried out using R version 4.3.2 (R Core Team 2023). Code used for the climate similarity analysis is available on GitHub: https://github.com/charles-cunningham/translocationClimate.

Four additional potential sites for introduction (sites 7-10 in Table S1) have also been identified in the Dart Valley in Devon (Simon Roper, *personal communication*). The climate analyses were repeated for these locations, using the same approach as for the West Sussex sites.

## RESULTS

### A. Community interactions

#### Natural enemies

Studies of parasitoids in the region under consideration are provided in Table 1. Confusingly, taxonomic revisions mean that few of the published names of *A. crataegi* parasitoids correspond to their current names. While pathogens have been mentioned, their identification is somewhat uncertain (nearly all accounts are historical; prior to the development of newer molecular methods) and most records are from outside our focal area (i.e., in Asia).

**Table 1.** Natural enemies.

| **Table 1. Natural enemies** |  |
| --- | --- |
| The Netherlands and Germany: *Cotesia glomerata* (sourced from the Netherlands) listed as a parasitoid of *A. crataegi* (sourced from Germany). *Cotesia glomerata* preferred *Aporia crataegi*-infested hawthorn leaves over uninfested hawthorn. | Geervliet, Vet & Dicke (1996) |
| Germany (Rhineland): *Apanteles glomeratus* parasitism reached a maximum of ~20-30% in 1955, and ~60-80% in 1956. Some 60-75% of the *Apanteles* cocoons failed to give rise to adults, mainly owing to (hyper)parasitism. | Wilbert (1959) |
| Germany (Rhineland): Parasitised by generalist *Apanteles glomeratus* and *A. pieridis*. Parasitism was variable and ranged from 0 to 28% | Wilbert (1960) |
| Germany: Parasitoids *Apanteles glomeratus* and *A. pieridis* caused high mortality in some places; *Pimpla instigator* and *Apechthis compunctor* parasitized up to 7 and 12% of the pupae, respectively, in 1956. | Blunck & Wilbert (1962) |
| France, South-East: Film of *Cotesia glomerata* parasitising *A. crataegi*. | Kan-van Limburg Stirum & Kan-van Limburg Stirum (2012) |
| Italy: Parasitoids recorded were *Apanteles difficilis*, *Pimpla instigator*, *A. spurius*, *Brachymeria scirropoda*, *Tricholyga segregata*, *A. glomeratus* and *Pteromalus puparum*. Populations reduced by polyhedral virus. | Martelli (1931) |
| Serbia: *Ceromasia rubrifrons* reared from pupa. | Stanković et al. (2014) |
| Unknown location, literature review: generalist *Apanteles glomeratus* identified as parasitoid. | Laing & Levin (1982) |
| Unknown location: cited report that *Bacillus thuringiensis* is highly virulent in *Aporia crataegi*. | Steinhaus (1951) |

The most frequently named parasitoid is *Cotesia glomerata* (=*Apanteles glomeratus*), which commonly attacks the large white butterfly, *Pieris brassicae*, and is already widely established in Britain. *Cotesia* (=*Apanteles*) *pieridis* is regarded as a synonym of *C. glomerata* (Broad, Shaw & Godfray 2016). *Pimpla instigator* is a synonym of *Pimpla rufipes*, a major host of which is, again, *P. brassicae*; this parasitoid is widespread in the UK. *Apechthis compunctor* is a Lepidopteran parasitoid that already occurs with scattered records from Cornwall to Norfolk, and north to North Wales and Yorkshire. *Apanteles difficilis* corresponds to *Cotesia cajae* (=*Cotesia perspicua*, =*Cotesia ofella*), all of which are reported already from England (Broad, Shaw & Godfray 2016). *Apanteles spurius* corresponds to *Cotesia spuria*, which is known already from England, Scotland, Wales and the Isle of Man (Broad, Shaw & Godfray 2016). *Brachymeria scirropoda* is a synonym of *Brachymeria tibialis*, which also parasitises a range of Lepidoptera and occurs in England. *Tricholyga segregata* (=*Exorista segregata*) is apparently *Exorista fasciata*, a polyphagous tachinid parasitoid of Lepidoptera (Tschorsnig, 2017) which occurs in England, Wales, Scotland and Ireland. *Pteromalus puparum* is a pupal parasitoid of Pieridae and Papilionidae butterflies, found in England, Wales and Scotland. In conclusion, all of the known parasitoids of *Aporia crataegi* in the regions of Europe under consideration already occur widely in Britain.

Other natural enemies (viral, bacterial and other pathogens) are also most likely to be shared with other species in the Pieridae butterfly family (the whites and yellows), to which *A. crataegi* belongs. The native resident British species in this family are the large white *Pieris brassicae*, the small white *P. rapae*, the green-veined white *P. napi*, the orange tip *Anthocharis cardamines*, the wood white *Leptidea sinapis*, and the brimstone *Gonepteryx rhamni*; with the clouded yellow *Colias croceus* visiting annually. GBIF records show that all of these species co-exist widely with *A. crataegi* across western Europe, making it unlikely that disease or parasitoid-mediated interactions between *A. crataegi* and the other pierids would have negative impacts on their distributions. These other pierid species co-exist with *A. crataegi* in both Brittany/Normandy and Catalonia (potential source regions for stock to release; see below).

Three of these pierid butterflies are migrants (Chowdhury *et al*. 2021), with *C. croceus* a ‘true migrant’ (Williams 1935, 1951), albeit with occasional overwintering, and *P. brassicae* and *P. rapae* as partial migrants (Williams 1935, 1951; Lack & Lack 1951; Baker 1969; Gilbert & Raworth 2005). Potentially migratory displacement may sometimes also occur in *P. napi* and *A. cardamines* (Baker 1969). Although the exact numbers of individual immigrants of each species of butterfly into Britain is unknown, trillions of insects immigrate to Britain from continental Europe annually (Hu *et al*. 2016). A study of migration in *one* Pyrenean pass estimated that “Of the butterflies, the most abundant family were the Pieridae (*Colias croceus*, *Pieris rapae* and *P. brassicae*) at 55,000 per year” (Hawkes *et al*. 2023). Some of the *Colias*, in particular, likely migrate directly from Catalonia, just south of the Pyrenees, to Britain. Thus, adult butterflies of related species immigrate into Britain in large numbers (probably many thousands annually), making it unlikely that the importation of *A. crataegi* from western Europe would bring new pathogens that have not previously arrived via the ongoing migrations of related species.

Pathogens transferred to larvae from host plants should also be considered. Commercial import numbers are typically treated confidentially, but an investigation established that Ireland imported at least a million living *Crataegus* plants from elsewhere in the EU in 2023 (Ryan 2023). For Great Britain, *Crataegus* was one of the most frequently imported trees / hedging plants imported from 2003 to 2013, comparable to *Betula*, for which over two million plants were imported during this period (Whittet *et al*. 2016). Import rates of *Crataegus* to the UK (not distinguishing Northern Ireland and Great Britain) have been ∼450,000 for bare root plants (mostly overwinter) and, more significantly, ∼210,000 for rooted plants in pots, including autumn and spring imports when plants are in leaf (EFSA 2023). Although imports are substantially reduced since Brexit (and since ash dieback), millions of *Crataegus* plants (and many *Prunus spinosa*) have been imported to Britain from continental Europe over the last three to four decades. Any pathogens linked to the host plants of *A. crataegi* are likely to have been imported repeatedly in the past (no importation of new plant material from continental Europe would be undertaken in any reestablishment project for this butterfly species). Moreover, the importation for horticulture of a wide range of potential host plants of *Pieris* species (plants in the Brassicales, including nasturtiums) has also occurred, plus the massive-scale imports of cabbages and other Brassicaceae vegetables. While individual insect pathogen transfer probabilities may be low, the magnitude and duration of this trade from continental Europe to Britain make it unlikely that plant-surface and plant-transferred pathogens exist in continental areas of western Europe without having arrived in Britain in the past.

#### Nectar sources

As with most butterfly species, selection of nectar plants is generally relatively flexible, but *Aporia crataegi* has an apparent preference for purple, red and pink flowers (Table 2). Several such species are widespread in the British countryside, including *Centaurea*, *Trifolium* and *Vicia* species. The availability of suitable nectar plants is not expected to be a constraint on the establishment of *A. crataegi* in Britain, although release sites with an abundance of nectar sources would be selected (mainly to avoid immediate dispersal away from them). *Aporia* would likely increase pollination services in the landscapes where it establishes; in Sweden, Lind *et al*. (2007) report it as one of the most important pollinators of the pyramidal orchid, *Anacamptis pyramidalis*, which also occurs on the chalk of the South Downs (see potential reestablishment sites, below).

**Table 2.** Nectar sources / adult feeding.

| Table 2. Nectar sources / adult feeding |  |
| --- | --- |
| France, Normandy: purple flowers, especially *Symphytum uplandicum* (comfrey), *Trifolium pratense* (red clover), *Centaurea nigra* (common knapweed) and *Vicia cracca* (tufted vetch). | Ratto (2008) |
| Sweden, Öland island: *A. crataegi* was one of two high frequency carriers of pollinia of the pyramidal orchid, *Anacamptis pyramidalis*. | Lind *et al.* (2007) |
| Slovenia: Nine plant species used for nectar, but *Knautia illyrica* (an Adriatic scabious) and *Vicia* aggr. *cracca* (tufted vetch) accounted for 80% and 84% of feeding occasions for males and females, respectively. | Jugovic, Crne. & Luznik (2017) |

#### Oviposition and larval host plants

We tabulated oviposition and larval host plant reports that are direct observations appropriate to a given location, rather than broader compilations of host records (e.g., as represented in field guides that lack information on frequencies of use, and that may not be geographically relevant). All recent oviposition host plant and larval records that are based on specific ecological field studies of *A. crataegi* in Belgium, northern France and Spain (the regions under consideration to source stock, see below) are from hawthorn *Crataegus monogyna* and blackthorn *Prunus spinosa* (Table 3). The only exception was a report that *Rosa* species may be used rarely in the mountains of Central Spain, but larvae from egg batches experimentally transferred to *Rosa* in that region failed to survive (Merrill *et al*. 2008).

**Table 3.**
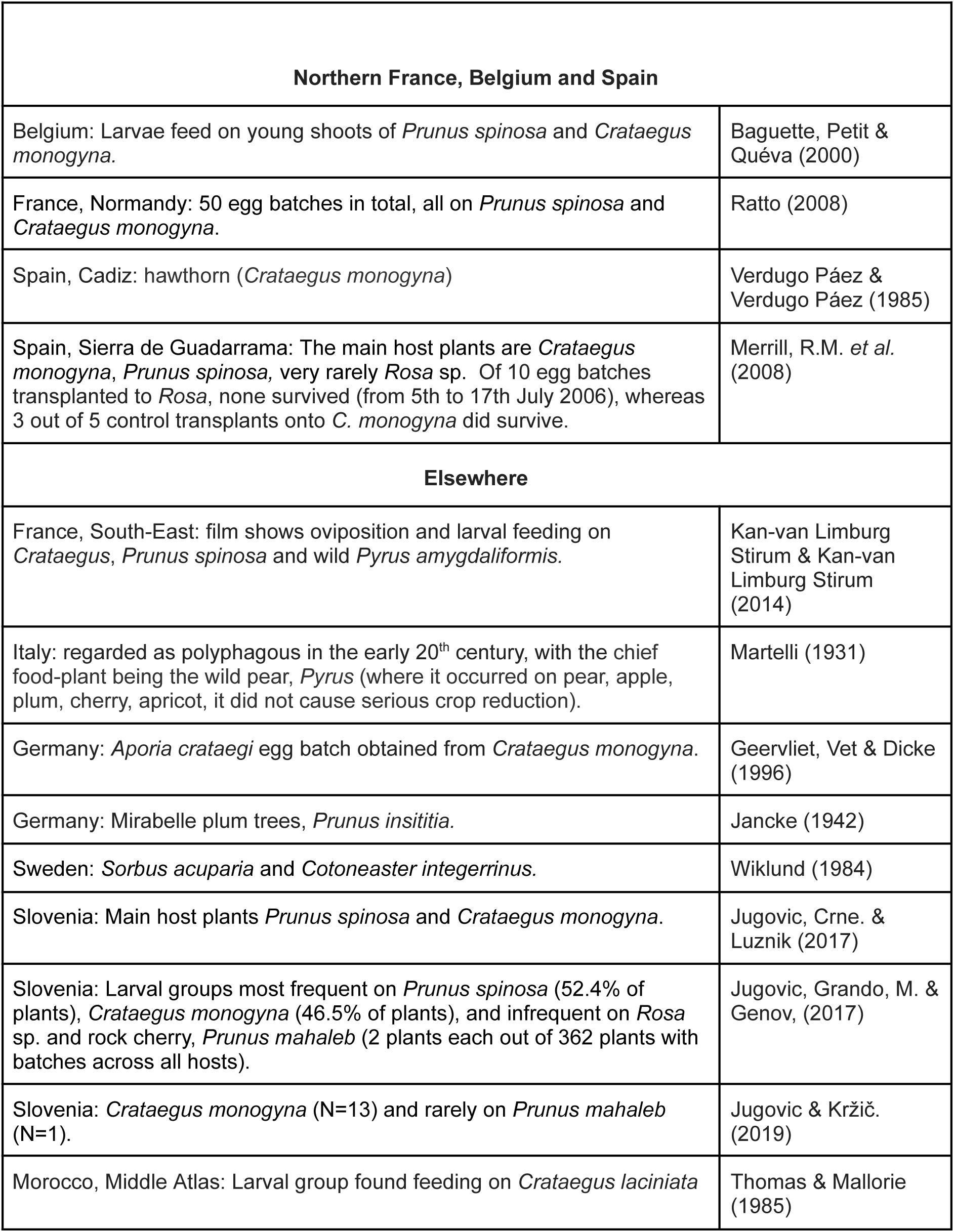
Oviposition and larval host plants.

| **Table 3. Oviposition and larval host plants** |  |
| --- | --- |
| **Northern France, Belgium and Spain** |  |
| Belgium: Larvae feed on young shoots of *Prunus spinosa* and *Crataegus monogyna*. | Baguette, Petit & Quéva (2000) |
| France, Normandy: 50 egg batches in total, all on *Prunus spinosa* and *Crataegus monogyna*. | Ratto (2008) |
| Spain, Cadiz: hawthorn (*Crataegus monogyna*) | Verdugo Páez & Verdugo Páez (1985) |
| Spain, Sierra de Guadarrama: The main host plants are *Crataegus monogyna*, *Prunus spinosa*, very rarely *Rosa* sp. Of 10 egg batches transplanted to *Rosa*, none survived (from 5th to 17th July 2006), whereas 3 out of 5 control transplants onto *C. monogyna* did survive. | Merrill, R.M. *et al.* (2008) |
| **Elsewhere** |  |
| France, South-East: film shows oviposition and larval feeding on *Crataegus*, *Prunus spinosa* and wild *Pyrus amygdaliformis*. | Kan-van Limburg Stirum & Kan-van Limburg Stirum (2014) |
| Italy: regarded as polyphagous in the early 20<sup>th</sup> century, with the chief food-plant being the wild pear, *Pyrus* (where it occurred on pear, apple, plum, cherry, apricot, it did not cause serious crop reduction). | Martelli (1931) |
| Germany: *Aporia crataegi* egg batch obtained from *Crataegus monogyna*. | Geervliet, Vet & Dicke (1996) |
| Germany: Mirabelle plum trees, *Prunus insititia*. | Jancke (1942) |
| Sweden: *Sorbus acuparia* and *Cotoneaster integerrinus*. | Wiklund (1984) |
| Slovenia: Main host plants *Prunus spinosa* and *Crataegus monogyna*. | Jugovic, Crne. & Luznik (2017) |
| Slovenia: Larval groups most frequent on *Prunus spinosa* (52.4% of plants), *Crataegus monogyna* (46.5% of plants), and infrequent on *Rosa* sp. and rock cherry, *Prunus mahaleb* (2 plants each out of 362 plants with batches across all hosts). | Jugovic, Grando, M. & Genov, (2017) |
| Slovenia: *Crataegus monogyna* (N=13) and rarely on *Prunus mahaleb* (N=1). | Jugovic & Kržič. (2019) |
| Morocco, Middle Atlas: Larval group found feeding on *Crataegus laciniata* | Thomas & Mallorie (1985) |

Earlier historical records (Pratt, 1983) and records further eastwards indicate that a wider range of host plants in the Rosaceae are used, especially other *Prunus* species (including orchard species such as plums; Jancke 1942), *Malus*, *Sorbus* and *Pyrus*. Records from south-eastern France confirm the use of *Crataegus* and *Prunus spinosa* but wild *Pyrus amygdaliformis* is also used (Kan-van Limburg Stirum & Kan-van Limburg Stirum 2014). Martelli (1931) reports wild pears as the main host plant in Italy, but also notes that *A. crataegi* has not caused “serious injury” to orchard trees. Genetically, the Italian *A. crataegi* belong to a different phylogenetic branch to those in Belgium, France, and northern Spain (and Slovenia, which also mainly uses *Crataegus monogyna* and *Prunus spinosa*), with the extinct British specimens also belonging to the Belgium-France-Spain clade (Todisco *et al*. 2020).

This suggests that *A.crataegi* has not represented a pest of orchards, at least for many decades, in the regions from which source material might be sought. For example, studies of insect pests of apple orchards in the Rhône valley in France do not mention *A. crataegi* (Simon *et al*. 2011); nor does a review of integrated crop management and organic systems for apple production in Europe as a whole (Tresnik & Parente 2007). There is no mention, either, of *A. crataegi* in three companion papers on the control of pests of apples and pears in northern and central Europe (Cross *et al*. 1999a, 1999b; Solomon *et al*. 2000), or in a more recent study of the control of invertebrate pests of pears across Europe (Shaw, Nagy & Fountain 2021). Nor could we find any recent mention of the butterfly as a pest of cultivated *Prunus*, cherries or plums (e.g., Jaastad *et al*. 2004; Quero-García *et al*. 2017). Therefore, we conclude that under modern horticultural conditions, including organic production systems, *Aporia crataegi* is not a pest of orchards and fruit production in northern or western Europe.

### B. Habitat assessment

#### Habitat and growth form of host plants - literature

Habitat assessments suggest that a wide range of habitats can be used by *Aporia crataegi*, ranging from dry grasslands through hedgerows and abandoned grasslands to woodland edges and rides (Table 4). In each case, habitats are characterised by scattered, and typically small, host plants. These may be relatively isolated host plants (of *C. monogyna* in particular) and suckering stems (especially of *P. spinosa*) in areas with scattered scrub, along hedgerows or at woodland edges. In some regions, host plants growing in sheltered conditions are especially favoured. This is likely to be the case in Britain, where summers are relatively cool. Shelter is not normally a major limitation because most areas with scattered scrub and overgrown hedgerows provide at least a degree of shelter.

**Table 4.** Habitat / growth form of host plants.

| **Table 4. Habitat / growth form of host plants** |  |
| --- | --- |
| Belgium: larvae feed on young shoots of *Prunus spinosa* and *Crataegus monogyna* growing at the margins of chalk grasslands. | Baguette, Petit & Quéva (2000) |
| France, Normandy: Meadows and hedgerows where 44 out of 50 egg batches were laid on plants under 2.5 m high; especially relatively isolated plants in shelter/sun. | Ratto (2008) |
| Sweden, Öland island: Various habitats including woodland / grassland / alvar steppe. | Lind *et al.* (2007) |
| Spain, Sierra de Guadarrama: Grassland, scrub and woodland sites, with a positive effect of host plant density on *A. crataegi* occurrence. Eggs were laid on the south side of host plants at high elevation, and on the shady northern side at lower elevations. | Merrill *et al.* (2008) |
| Slovenia: Dry karst meadows with suitable nectar plants, and *Prunus spinosa* and *Crataegus monogyna* larval hosts. | Jugovic, Crne. & Luznik (2017) |
| Slovenia: Karst meadows. Groups of larvae found on smaller host plant individuals, in particular microclimates. Larval nest densities were positively but not significantly correlated with host plant density. | Jugovic, Grando & Genov (2017) |
| Slovenia: Karst meadows with hedgerows consisting mainly of *Crataegus monogyna*, *Prunus spinosa*, *Prunus mahaleb*, and *Rosa* sp. Eggs were laid on the upper side of the leaves on relatively small (low, small diameter) host plants. | Jugovic & Kržič (2019) |

A range of events has increased the availability of such habitats over the past 70 years: myxomatosis killed off rabbit populations from the mid 1950s, thereby generating substantial amounts of hawthorn scrub (Figures 3, 4) and blackthorn suckering. Reduced frequencies of hedgerow management (cutting) have accompanied agricultural field margin protection policies; and additional scrublands have developed as a consequence of recent rewilding projects (Figure 2), as well as connectivity and similar conservation schemes (e.g., extensive grazing that prevents succession to denser scrub or woodland).

**Figure 2.**
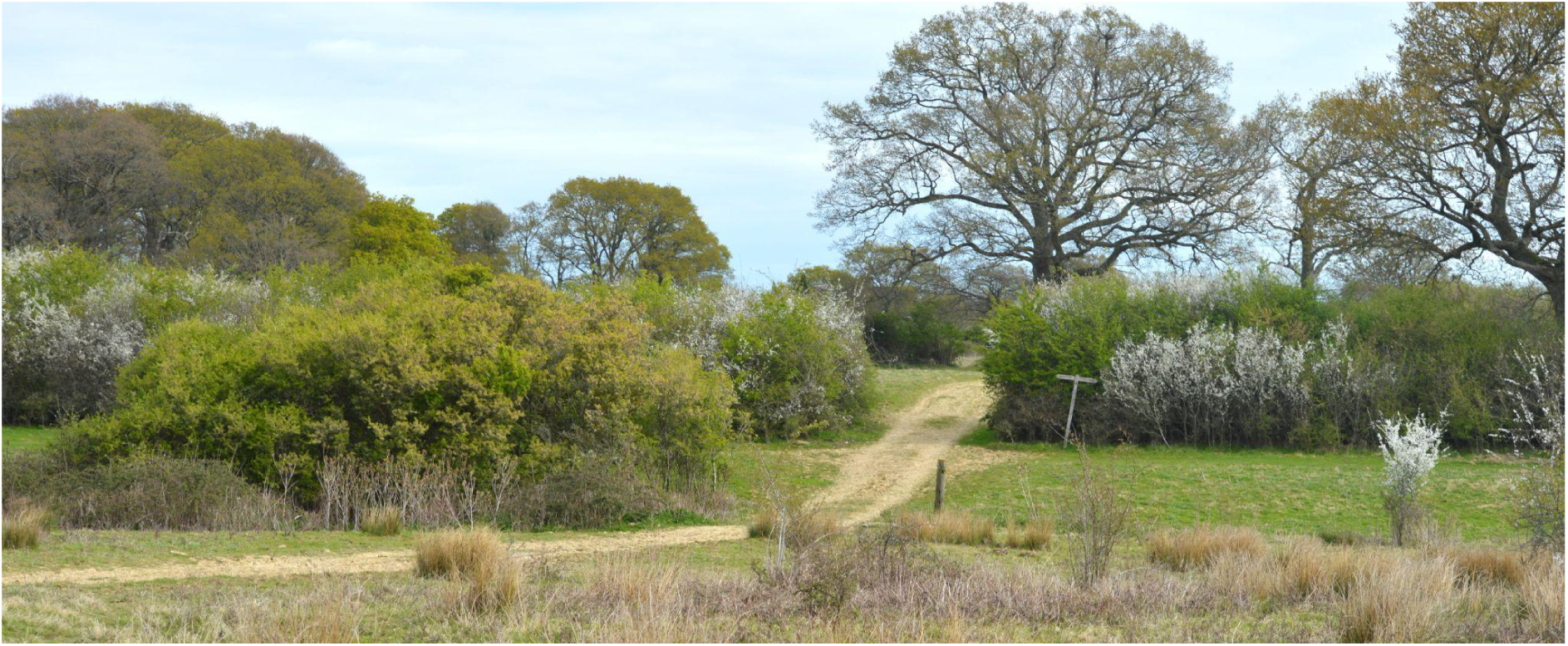
Overgrown hedgerows and scattered white-flowering *Prunus spinosa*, with fresh green growth of *Crataegus monogyna*, at Knepp Wildlands (photo © Chris Thomas)

**Figure 3.**
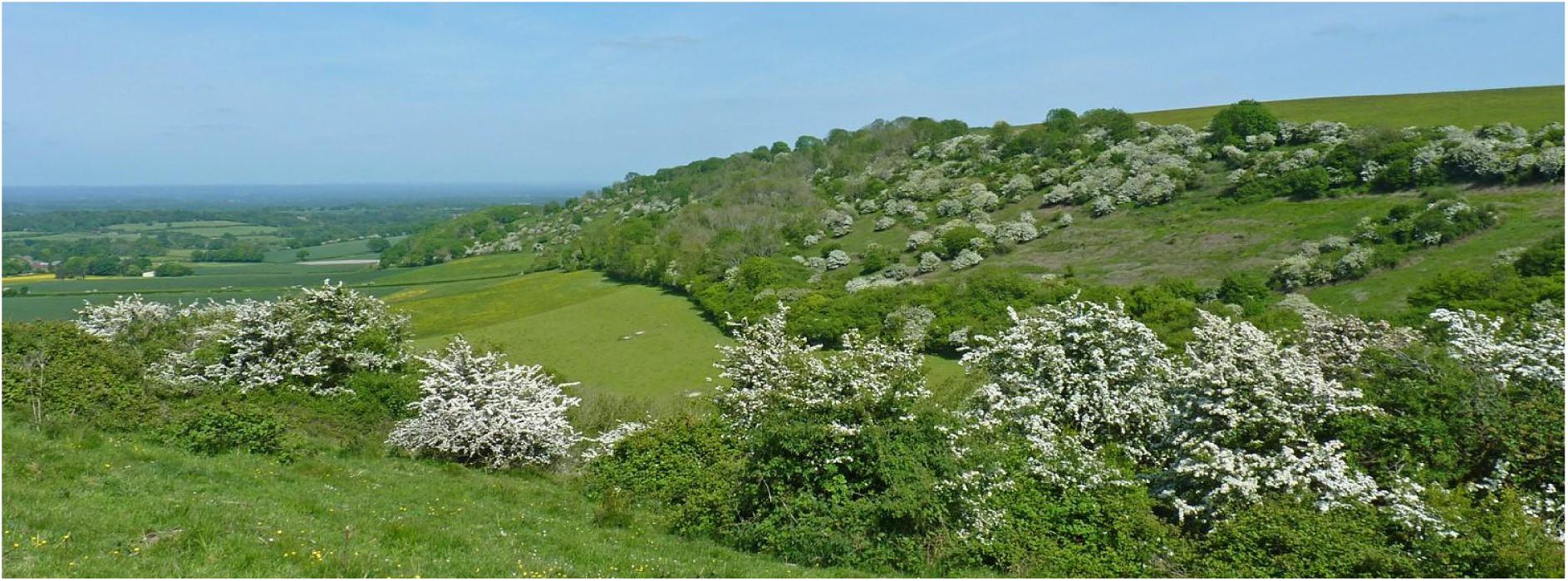
Scattered flowering *Crataegus monogyna* on the South Downs escarpment of West Sussex; view from Sullington Hill towards Barnsfarm Hill (photo © Neil Hulme)

**Figure 4.**
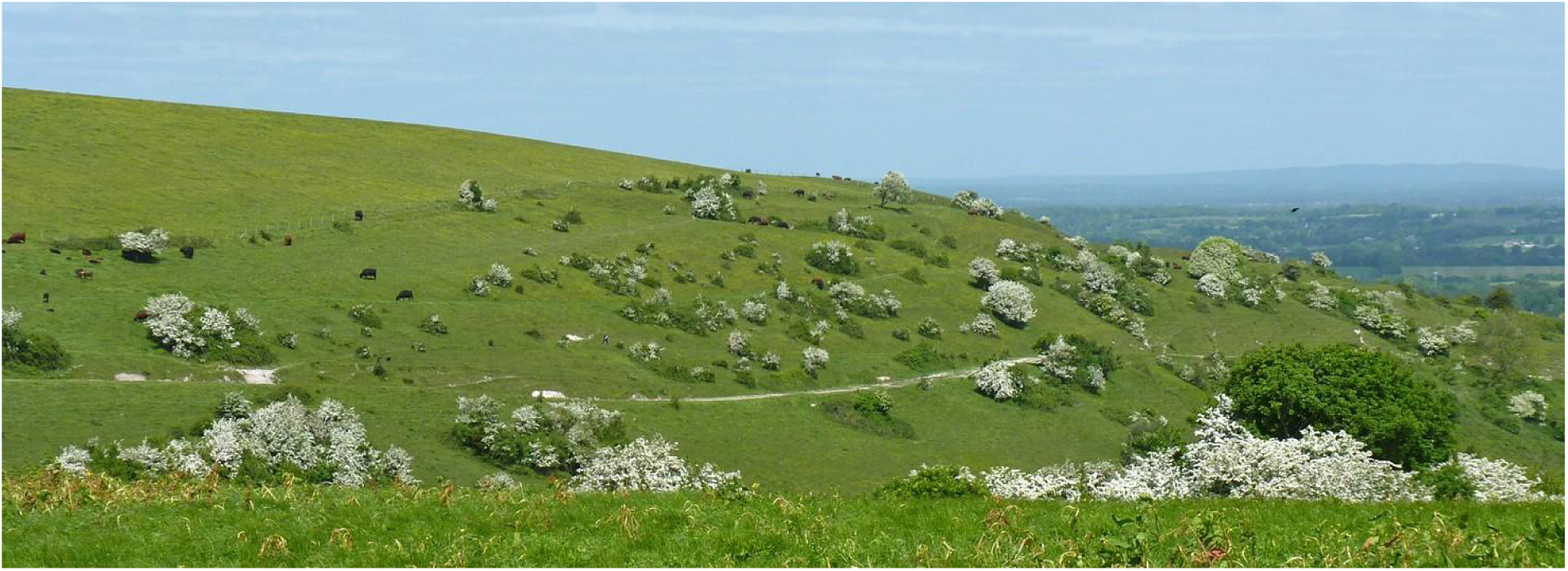
Sullington Hill, West Sussex, with flowering bushes of *Crataegus monogyna* (photo © Neil Hulme)

#### Habitat at potential sites for establishment

Based on prior experience and fieldwork in West Sussex, England, the following sites were identified as meeting habitat criteria, based on their host plant, growth form, nectar sources, shelter and landscape attributes (Table 5). They all fall within an ∼11 km by 9 km landscape, which is expected to operate as a patchy-population or metapopulation on a multi-year time scale (see Dispersal, below).

**Table 5.**
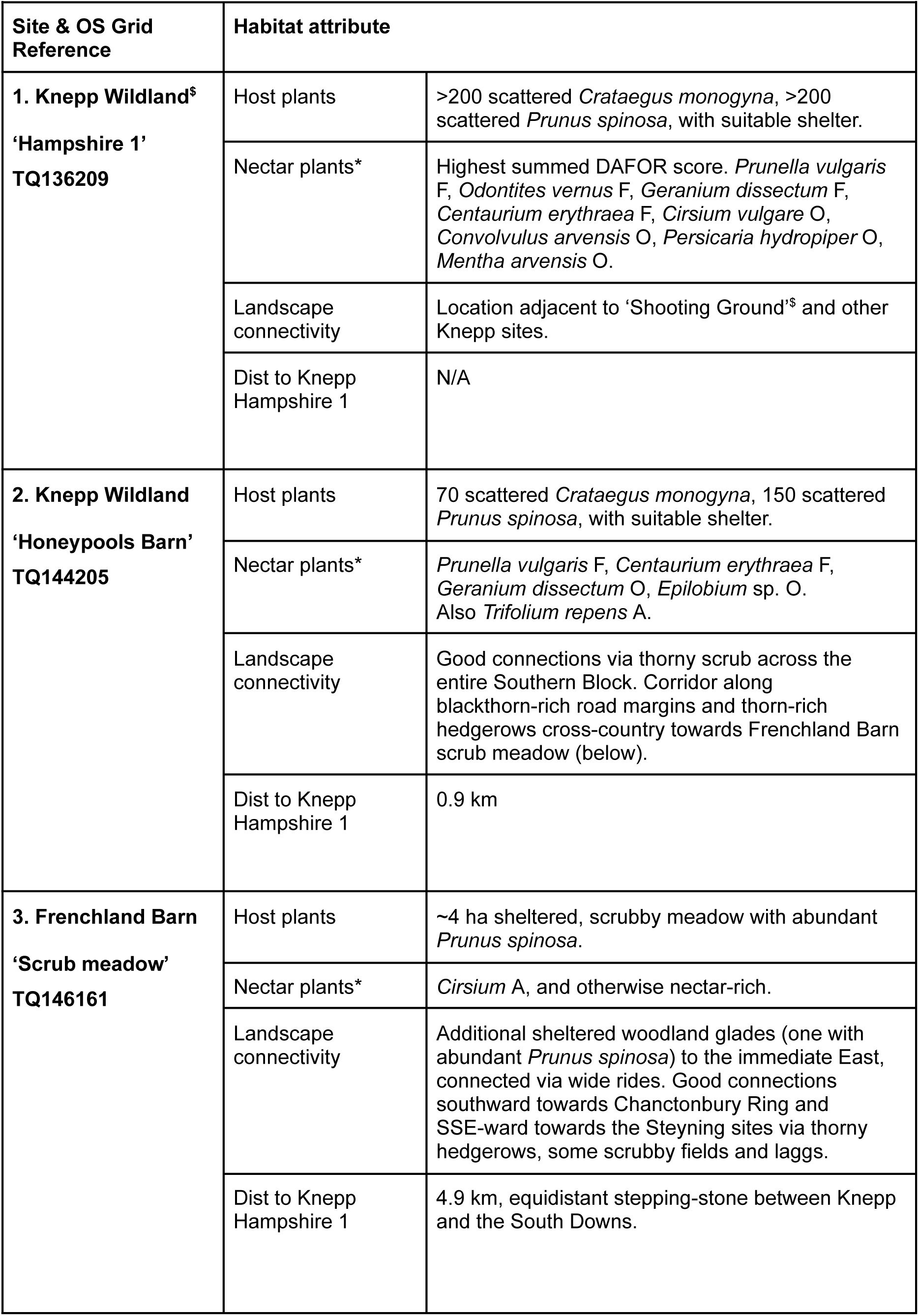
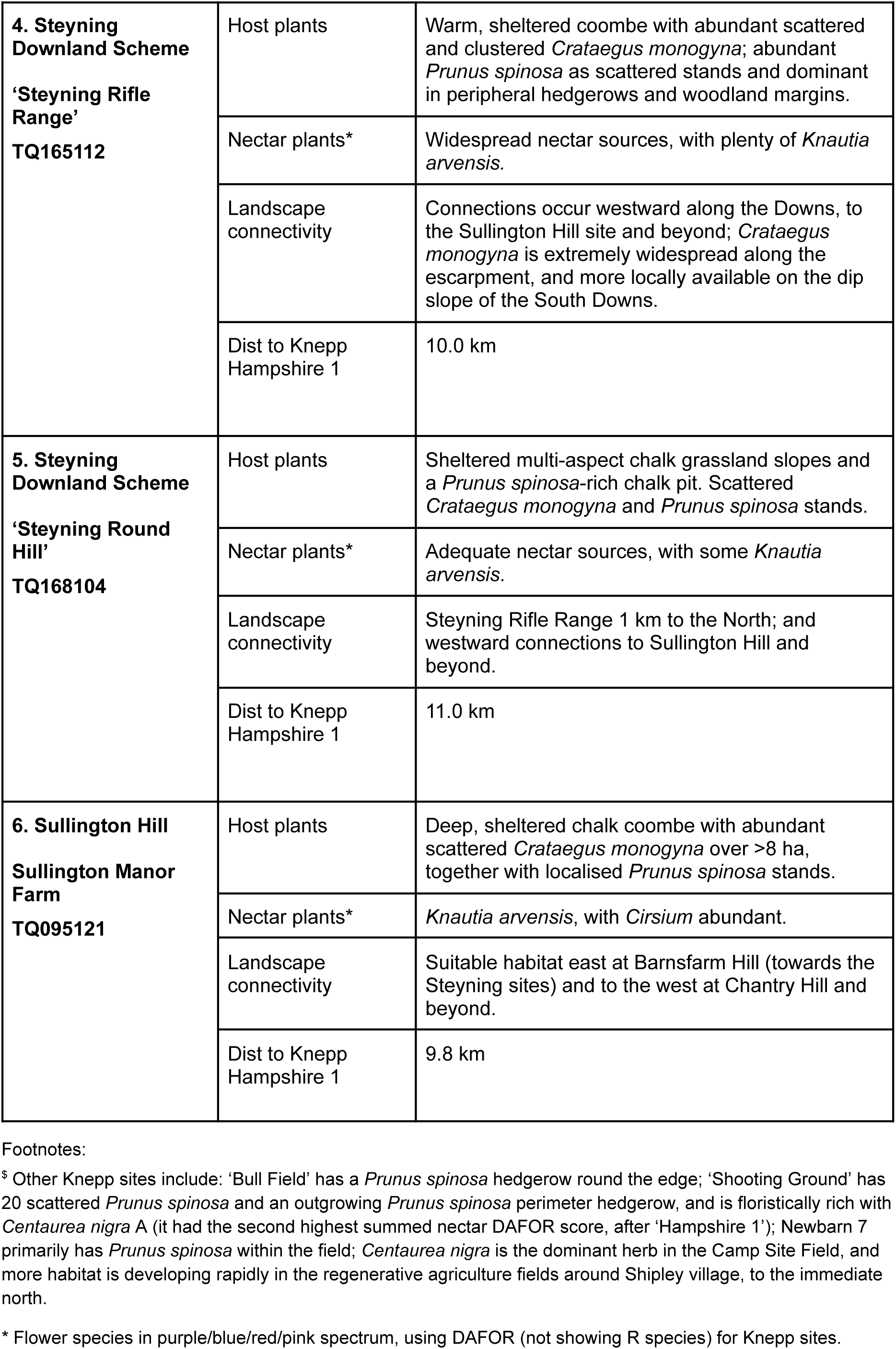
Potential initial reestablishment sites in West Sussex, England.

These include regenerating scrubland within the Knepp Wildland project area (Figure 2), which has extensive grazing by cattle, ponies, pigs and wild ungulates; Frenchland Barn, containing woodland management that provides openings within woodland and rides; and chalk sites in the South Downs, mainly characterised by extensive grazing management (Figures 3, 4). Many other locations are potentially suitable within the same landscape.

Although the habitat needs of the species appear to be met in these and other habitat patches within the target landscape, there inevitably remain some areas of uncertainty. Therefore, the proposed release sites include different soil types, different mixtures of larval host plants and nectar sources, different browsing regimes by different herbivore species (from rabbits and sheep to deer and cattle) and different distances to nearby habitat patches. This variation will enable any initial experimental release programme to generate new knowledge that will inform priorities when identifying subsequent release sites and when considering management options.

### C. Climatic suitability assessment

#### Sensitivity of populations to climate

Several articles include *Aporia crataegi* within multi-species studies of butterfly responses to climate change, especially in the Mediterranean region (Table 6). The distribution of *A. crataegi* across Europe as a whole (Figure 1) suggests that it is largely a montane species in the south, but occurs in the lowlands further north. This implies that climate, and specifically temperature, is an important determinant of the species’ distribution. In the Spanish mountains, the butterfly is more abundant (per host plant) at higher elevations, but the upper host plant elevational limits likely constrain the butterfly’s capacity to retreat to higher elevations (Merrill *et al*. 2008); it has shifted phenology in response to climate warming (Goded *et al*. 2024).

**Table 6.** Climatic limits and responses.

|  |  |
| --- | --- |
| Spain, Sierra de Guadarrama (central Spain): The species is apparently limited by hot temperatures at low elevation, and larval survival increased with elevation. Absence of suitable host plants at higher elevation limits the capacity of *A. crataegi* to survive climate change by colonising higher elevations. | Merrill, R.M. *et al.* (2008) |
| Spain, Sierra de Guadarrama: *Aporia crataegi* did not show population declines at higher or lower elevations in central Spain. | Caro-Miralles & Gutiérrez (2023) |
| Spain: *Aporia crataegi* advanced their flight dates (emergence earlier in the year) in 2017-2022, compared to 1985-2005, but did not exhibit an increase in elevation; also in the mountains in central Spain. | Goded *et al.* (2024) |
| Spain, Catalonia: Periodic population crashes, likely related to unsuitable climatic conditions. | Ubach *et al.* (2022) |
| Greece: *Aporia crataegi* showed a substantial decline in abundance between 1998 and 2011/2012 in a Greek National Park. In contrast, low elevation butterfly species tended to increase during a period when the regional climate warmed by 0.95°C. | Zografou *et al.* (2014) |
| England / Europe: Climate / distribution modelling reveals that central, southern and eastern England are climatically suitable for *A. crataegi* and projected to be amongst the most suitable climatic areas for the butterfly in Europe, with declines projected in lowland / southern Europe (Figure 6). | Carroll *et al.* (2009) |

Merrill *et al*. (2008) concluded that *A. crataegi* was temperature-limited at low elevations in the Sierra de Guadarrama (central Spain), given that its *C. monogyna* and *P. spinosa* host plants remained common at lower elevations (where the butterfly was absent) and that it laid eggs on the ‘shady’ (cooler) side of host plant bushes at low elevations (where ambient temperatures are higher). Although *A. crataegi* has not declined in overall abundance or shifted elevation in this region in recent years (Caro-Miralles & Gutiérrez 2023; Goded *et al*. 2024), it has declined in Greece, where *A. crataegi* showed one of the steepest declines in numbers, in line with regional warming (Zografou *et al*. 2014). Together, these results suggest climate sensitivity in the Mediterranean region. It may be vulnerable to hotter climates in some regions, given the absence of higher elevation populations of host plants.

### Climate at the time of extinction from England

The Central England Temperature (CET) record (National Climate Information Centre 2024) is consistent with the hypothesis that the decline of *A. crataegi* was linked to low temperatures towards the end of the 19^th^ and in the early 20^th^ centuries. The late 1870s through to 1892 were particularly cold, with further cool spells in 1907 to 1909, and cold years in 1922 to 1924 (Figure 5), immediately prior to the butterfly’s final records, in 1923 or 1925. No single *year* has been as cold as the 1875-1925 *average* since the 1985 to 1987 period, and only one year (1962) has been as cold as 1924 (mean spring/summer temperature of 10.65 °C for both years), the year preceding the butterfly’s last possible sighting as a breeding species. The average daily spring and summer temperature of the period 1990 to 2023 inclusive is 12.65 °C, or 1.15 °C hotter per day than the 1875 to 1925 average. The average of the last seven years (2017-2023 inclusive) is 13.07 °C, some 1.57 °C warmer than the average for the period preceding the species’ extinction (National Climate Information Centre 2024).

**Figure 5.**
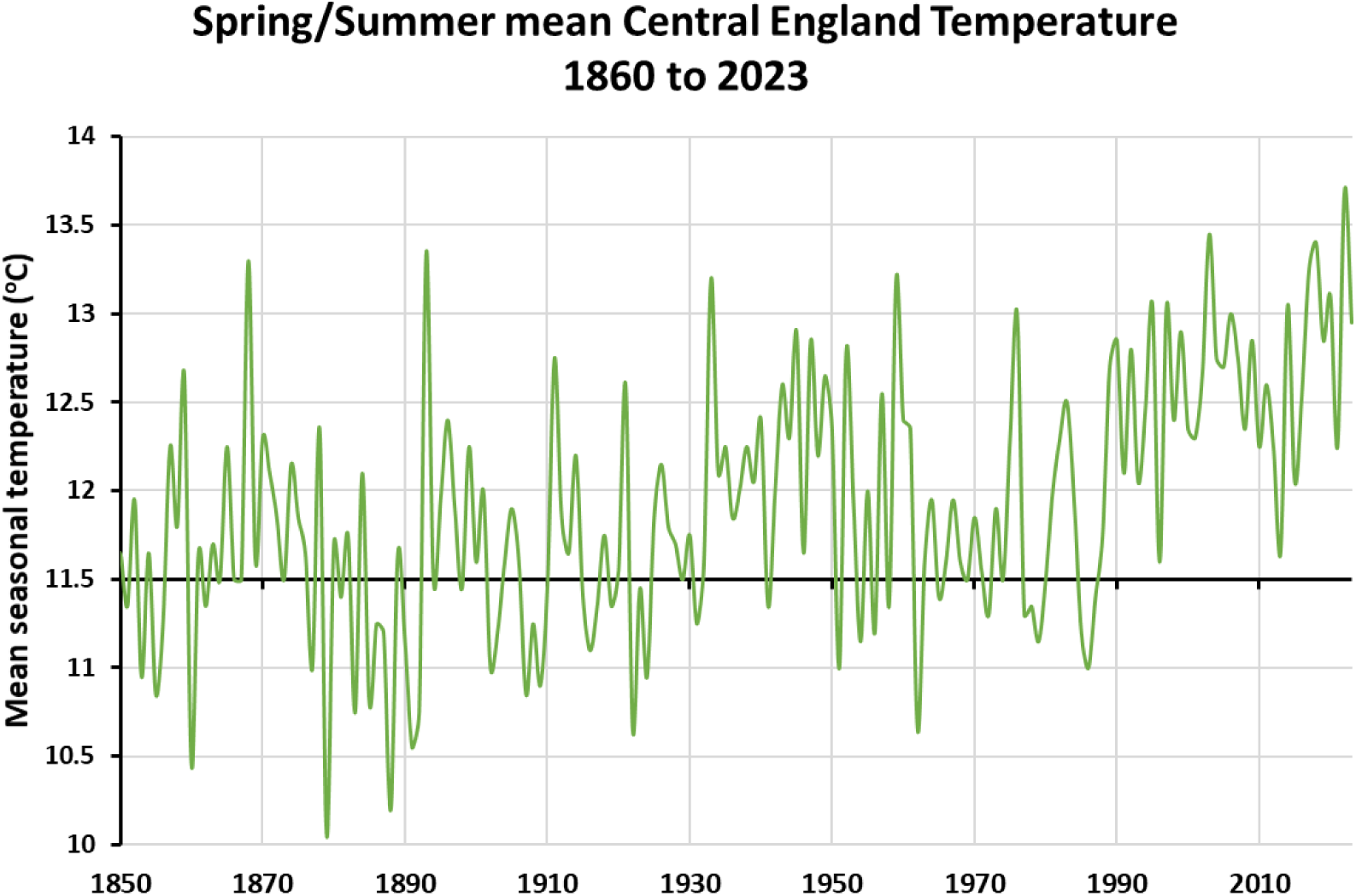
Central England Temperature (CET) average of seasonal daily temperatures for spring (March-May) and summer (June-August) for 1850 to 2023. The horizontal line shows the average daily temperature of spring and summer in the CET from 1875 to 1925 inclusive, at 11.5 °C.

This potentially translates into a substantial proportional increase in accumulated ‘development time’ (or growing degree days) above a thermal threshold of 5 °C to 10 °C (the normal range of minimum temperature thresholds for butterflies in northern Europe). Spring sunshine hours *in England* have also increased, by around 15% between the 1910s and 2010s, while summer sunshine hours have remained stable or increased very slightly (Met Office 2024). Increased spring sunshine hours facilitate the capacity of larvae to thermoregulate, further increasing the potential developmental rate of *Aporia crataegi* larvae under present-day climatic conditions.

#### Distributional potential

Carroll *et al*. (2009) modelled the recent distribution of the species across Europe in relation to climate variables - using Generalised Additive Models and Generalised Linear Models for three climate model scenarios for 2021 to 2050 (which all gave very similar results). An example is given in Figure 6, which shows that the climate in most of England (apart from the South-West and Lake District) is expected to be suitable under past (1961-1990), current and future climates (2021-2050); with a corresponding decline in southern Europe. These models and scenarios suggest that England has some of the most suitable climates in Europe for the species. However, the capacity of *A. crataegi* to expand its range to higher latitudes may be limited by dispersal (see below).

**Figure 6.**
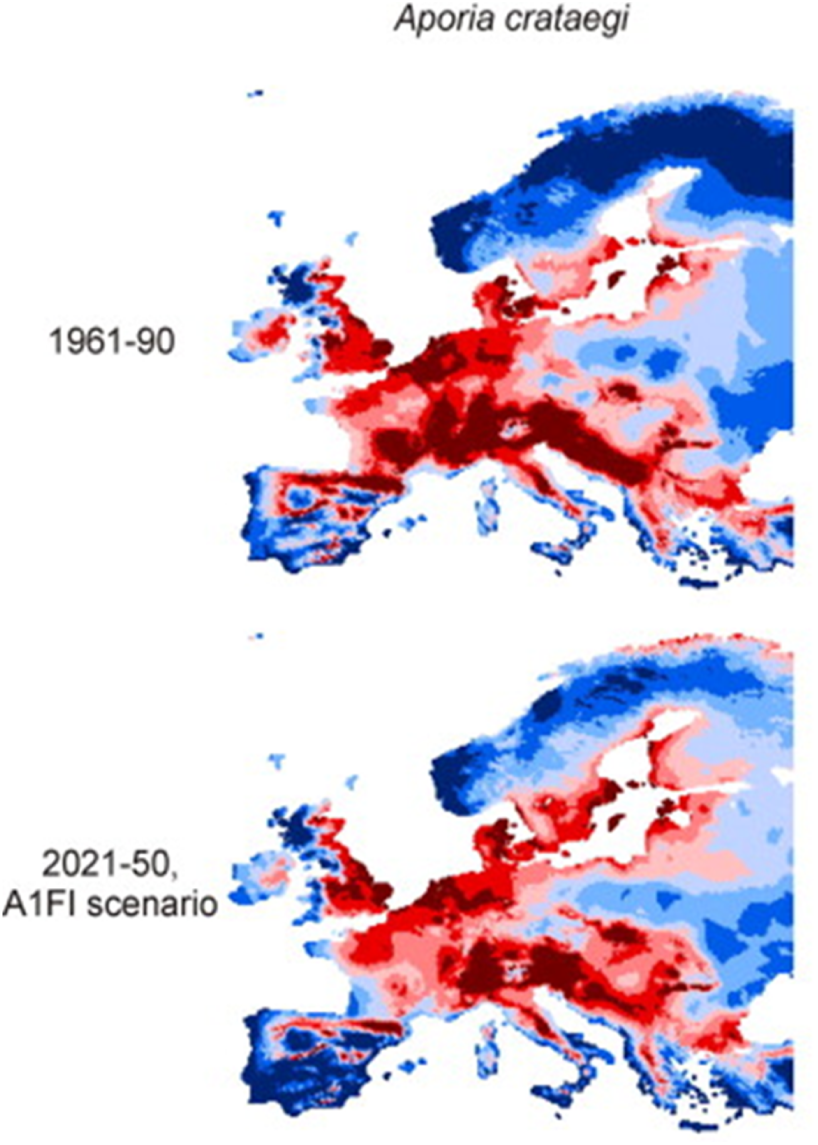
Modelled (GAM) climate suitability of Britain and Europe for *Aporia crataegi* for the late-20^th^ century and for a 2021 to 2050 climate scenario (selected panels from Fig. 2 in Carroll *et al*. 2009; © 2009 Elsevier Ltd.). Darker red indicates high climate suitability; blue climatically unsuitable areas.

#### Availability of source locations with climates similar to south-central England

Sourcing the most appropriate material to reestablish *A. crataegi* in Britain is usefully informed by the similarity of climatic conditions between proposed source populations and proposed reestablishment sites. Presented as climate surfaces for four climate variables, Figure 7 shows that southern England falls within the range of climatic conditions where current populations of the species (Figure 1) exist within continental Europe. Figure 8 shows the locations of the areas in Europe with the most similar recent climates to a key proposed establishment sites (red spot) in West Sussex, England, for each of the four climate variables considered separately. All four climate variables indicate climate similarity of West Sussex to areas of continental Europe that support populations of *Aporia crataegi* (Figure 1), as well as to other areas in southern, central and eastern England. For example, the suggestion that English winters might not be cold enough for this species is unfounded, given that populations survive in continental areas similar to the south-west of England, where the winter climate is warmer (as represented by the mean temperature of the coldest month - January), just as they occur further east in Europe, where the winters are colder (Figures 7, 8).

**Figure 7.**
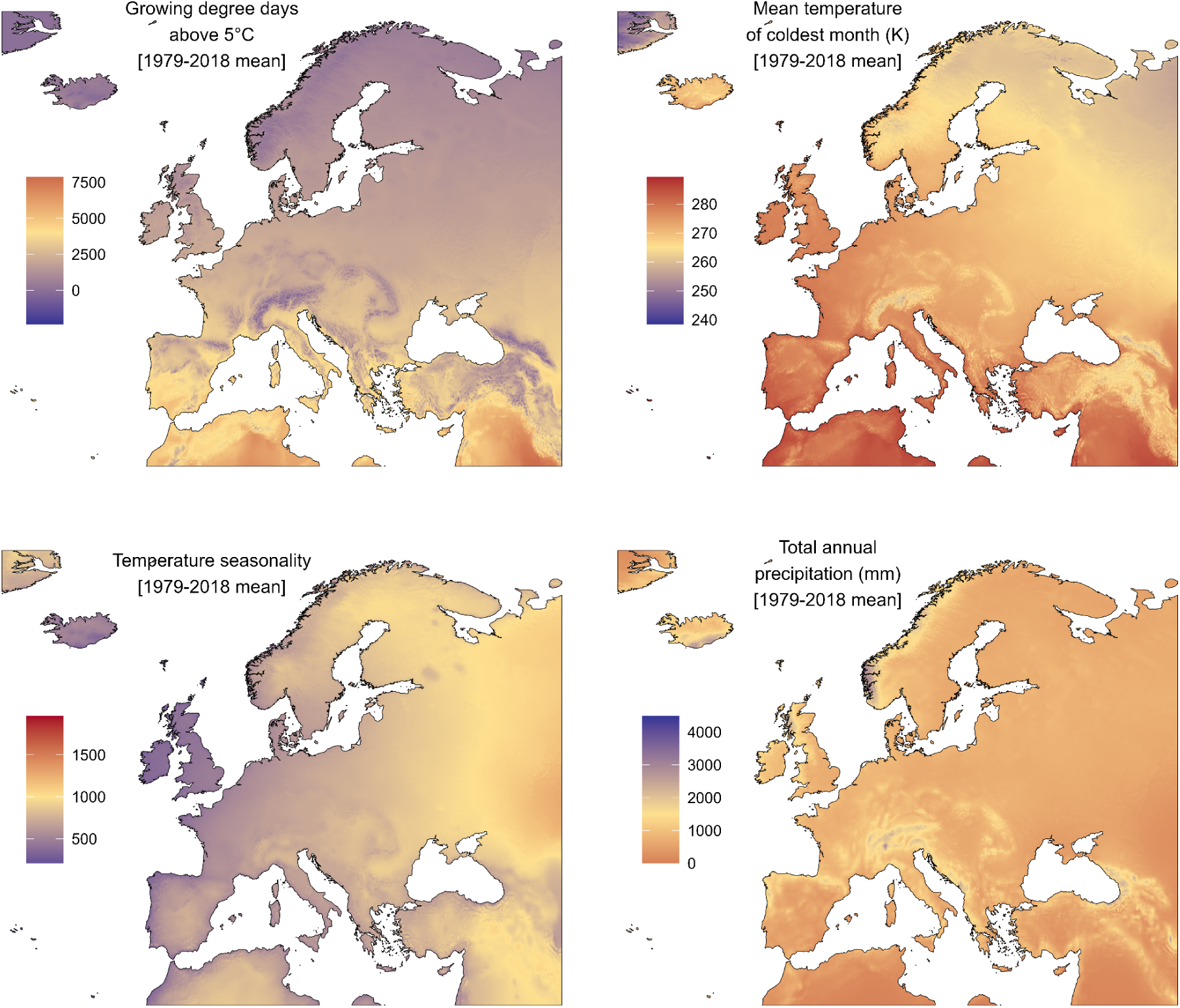
Distribution of four climate variables across Europe.

**Figure 8.**
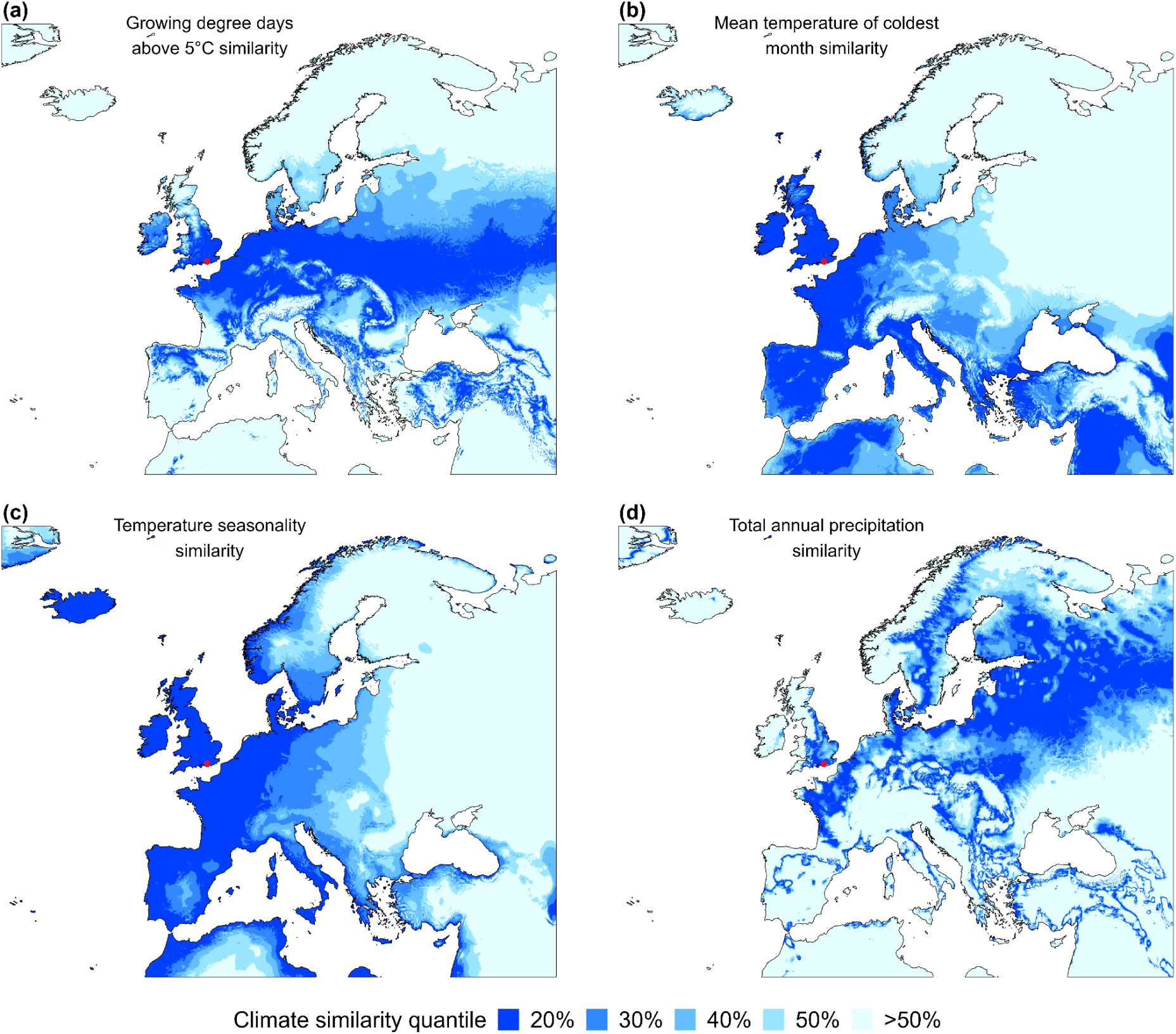
Similarity in quantiles of four climate variables to the West Sussex translocation sites (red dot), England.

Combining these variables (the overlap in the percentiles from the four panels of Figure 8) highlights the climatic similarity of northern France and localised mid-elevation areas in southern France and the Iberian Peninsula (Figure 9a). However, the main consideration is whether conditions are warm enough in lowland England, given that *A. crataegi* was hypothesised to die out in the UK following several cold decades, and that the species occurs in parts of Europe that are wetter as well as areas that are drier than lowland England. If we consider the three thermal variables (winter cold, growing degree days, thermal seasonality), similar areas are highlighted (Figure 9b). This provides a greater range of locations from which to source potentially suitable stock for reintroduction, albeit still with a focus on north-western Europe and Iberian Peninsula mountain ranges.

**Figure 9.**
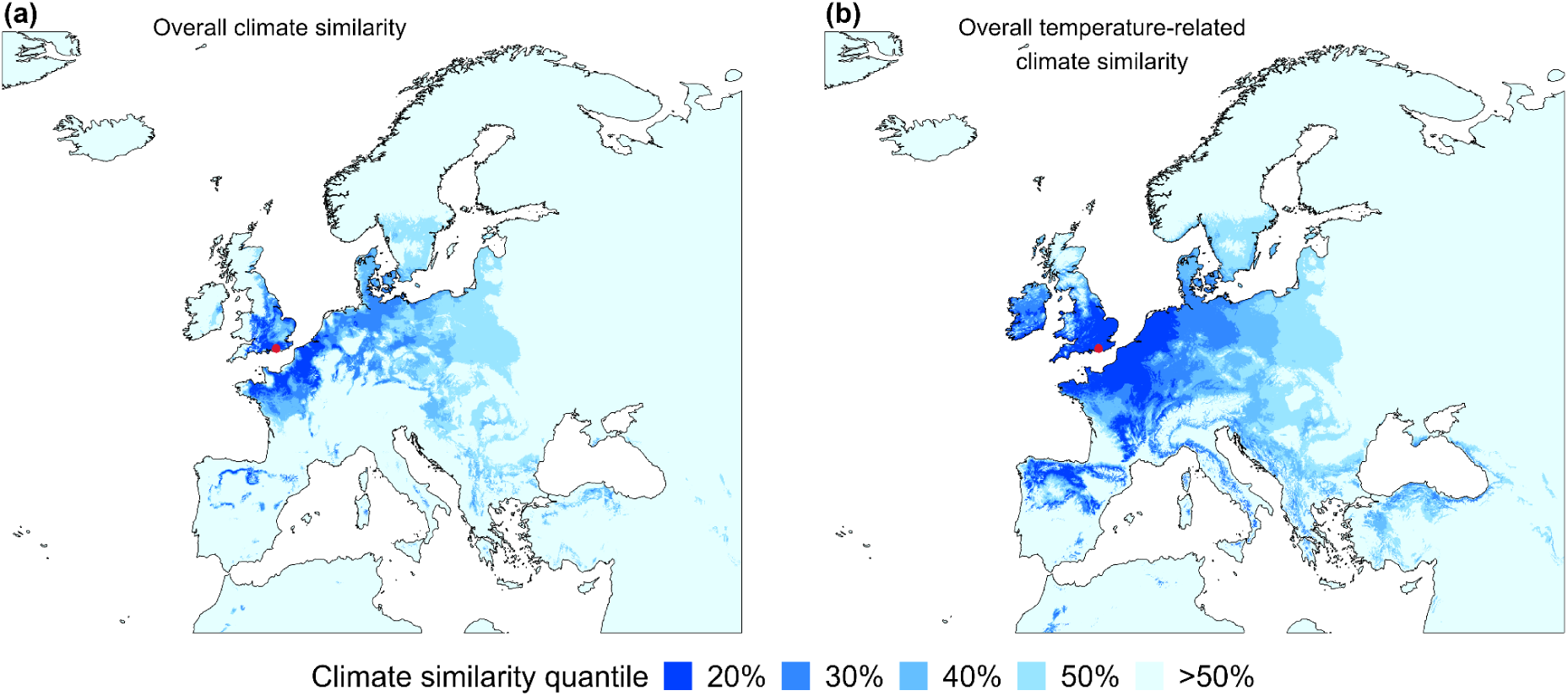
Overlap between the climatic match for all four climate variables considered (a), and for the three temperature-related variables (b), to the potential West Sussex translocation sites. 20% corresponds to locations in the top 20% similarity for all variables considered.

Climate is not the only determinant of the species’ distribution (which includes habitat, host plants, nectar plants; as outlined above), and hence it is important to identify more specific locations as potential sources of *A. crataegi* stock for reintroduction. Two key areas are recognised: parts of Brittany (and potentially Normandy) in France and mid-elevations in the eastern Pyrenees in Spain and France, and Massif Central in France, where concentrations of records of *A. crataegi* overlap with the climate-match areas (Figure 10), although other areas of western Europe also have matching climate, especially in France (Figure S1).

**Figure 10.**
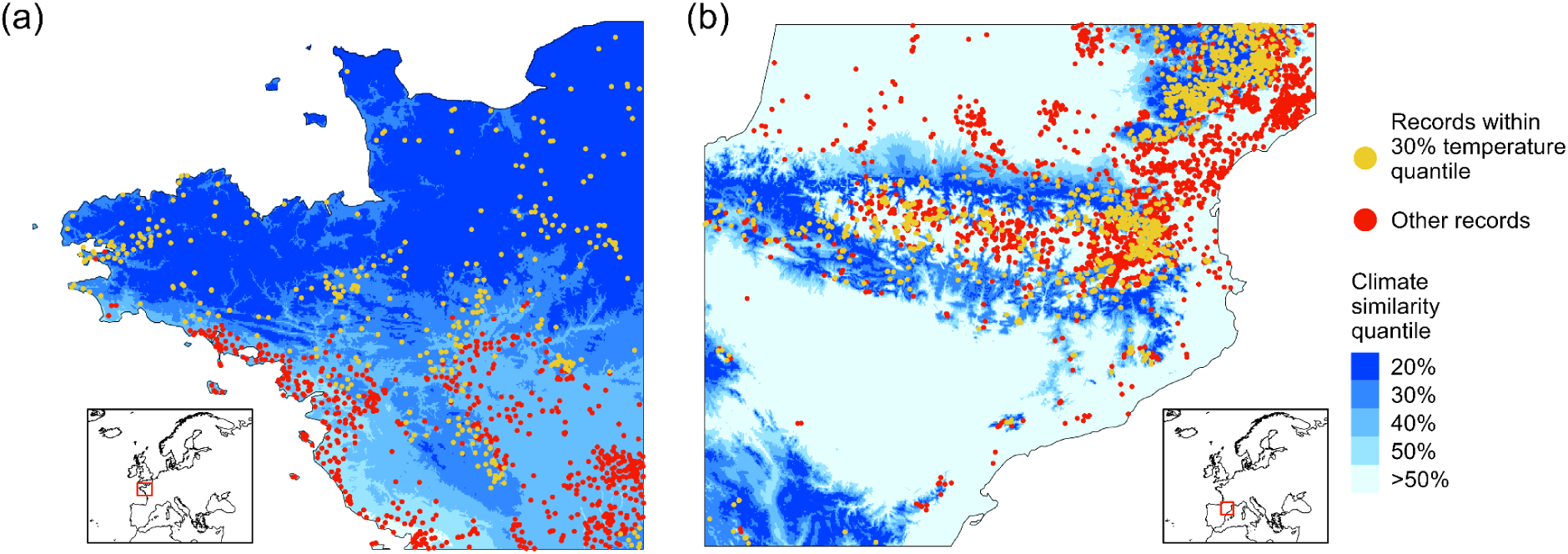
North-western France (a) and the Pyrenees (b), illustrating areas of overlap of distributional records of the species over the last decade (from GBIF) with the temperature match (30%) of the area from Figure 9b. Climate match to West Sussex sites: species records with similar (yellow) and different (red) climatic conditions are shown.

Repeating these analyses for the climate of additional potential British reintroduction sites in the Dart valley in Devon (sites 7-10 in Table S1) generated similar results. Locations where the species occurs at present in continental Europe are both more and less extreme for all climate variables considered (Figure 7), with the best climate similarities again in north and north-west France, parts of central France, and montane areas of the Pyrenees and other Iberian peninsula mountain ranges (Figures S2-S5). The similarity of the Devon and West Sussex results suggest that the same population sources would likely be suitable for releases in both British translocation areas.

### D. Dispersal

#### Movements of adult butterflies

The available evidence of butterfly movements is consistent across mark-release-recapture studies (Table 7). Despite being more mobile than some other butterflies, most movements have been of individuals recaptured within the same site where they were marked, and few individuals have been recorded as moving further than 1 km (maximum of 3.5 km). All studies report that males are considerably more mobile than females, providing genetic and some population connections across a landscape where habitat patches are scattered over 1 to 11 kilometres (see above). As such, a landscape approach to the reestablishment and conservation of this species is required.

**Table 7.** Dispersal studies.

|  |  |
| --- | --- |
| Belgium: 58% of recaptures were relatively short within-patch movements. Dispersal distances between patches fitted to inverse power law function (i.e., long-tailed) with a maximum recorded distance of 1.59 km. Higher immigration and emigration rates for the smallest patches. | Baguette, Petit & Quéva (2000) |
| France, Normandy: Males more mobile than females. Individuals moved between meadows. | Ratto (2008) |
| Sweden, Öland island: mean dispersal distances (over 2.5 days average) was 315 m for males and 182 m for females. Three out of 94 recaptures were over 1 km. | Lind *et al.* (2007) |
| Slovenia: Most recaptures were within sites, with some movement over a few kilometres, especially for males. Male median distance was 604 m, max 3.5 km; the only between-site movement for a female was 1.4 km, with mean daily movements of females only 13.3 m. | Jugovic, Crne, & Luznik (2017) |

Nonetheless, the frequencies of within-site recaptures and modest between-patch distances (only 3 out 94 recaptures were over 1 km in Lind *et al*. 2007), especially for females, imply that relatively separate breeding concentrations are likely to be maintained within suitable habitat patches and areas of the landscape. The rarity of dispersal distance over 1 km by females (the colonising gender), the scarcity of large *A. crataegi* populations close to the English Channel in France, Belgium and the Netherlands (due to intensive land use), and the need for colonists not only to cross the Channel but also to find suitable habitats once they arrive, make unassisted recolonisation of Britain unlikely in the short to medium term.

## DISCUSSION

There is considerable potential to reestablish the black-veined white butterfly, *Aporia crataegi*, in lowland England. The primary cause of extinction of the species in the early 20^th^ century is normally stated to have been the climate. However, the climate of lowland southern, central and eastern England today is considerably warmer than in the years preceding the species’ extinction from Britain, and is projected to be one the most climatically suitable regions of Europe in the coming decades (Figure 6). Furthermore, the proposed landscape for reestablishment in West Sussex, England, contains many sites that meet the habitat, host plant and nectar requirements of the butterfly (Table 5), at a spatial scale relevant to the dispersal and population connectivity of the species (Table 7). While success can never be guaranteed, all the potential indicators suggest that a substantial and extensive metapopulation could be established, from which subsequent relocations within Britain could then be coordinated.

Previous re-establishment attempts have almost certainly been too small scale, with too few adults released over too short a period of time, in too small a habitat patch. The most ambitious of these releases took place at Holmwood Common, just south of Dorking, Surrey, in the mid-1970s, where several hundred adults were released. This release coincided with the great drought of 1976 and the subsequent wet June of 1977 (Oates & Warren, 1990).

Thus, any release programme should be planned to take place over several years to avoid single-year climatic events that prevent establishment (and equally to take advantage of particular ‘good years’). Given the dispersal behaviour of *A. crataegi* (above), the likelihood of success will be higher if habitat area is scattered over substantial areas of the landscape, as in the ∼100 km^2^ Knepp to Downland landscape considered here. This overlaps with the Weald to Waves initiative area, coordinated by the Knepp Wildland Foundation, which aims to create a 160-kilometre nature recovery corridor across Sussex. Landowners and managers are being encouraged to increase scrubland habitat along the corridor, thereby benefiting any re-introduced black-veined whites as well as other threatened scrubland species.

### Co-benefits for other species

Species associated with scrubland, infrequently managed hedgerows and mosaics of early to later successional vegetation include several bird species that are red listed in the UK, including nightingales (*Luscinia megarhynchos*), cuckoos (*Cuculus canorus*) and turtle doves (*Streptopelia turtur*), and the regionally-extinct red-backed shrike (*Lanius collurio*), another species the Knepp Wildland Foundation is working with Natural England to reintroduce. There is a rich invertebrate fauna associated with blackthorn and hawthorn, some as foliage or even blossom obligates (such as the locally-distributed sloe pug moth *Pasiphila chloerata* and the sloe carpet moth *Aleucis distinctata*, the latter a southern and south-eastern rarity that occurs at Knepp). Most associated insect species utilise a range of shrubs, though they strongly favour these two plant species, such as the hawthorn shieldbug *Acanthosoma haemorrhoidale*. There is a sequence of invertebrates that utilise hawthorn and blackthorn in various stages of tree development, culminating with a scarce beetle that breeds in the trunks of veteran hawthorns (the hawthorn jewel beetle *Agrilus sinuatus*). Many of these species have declined significantly in recent decades, including the once-common lackey moth *Malacosoma neustria*. This project will help restore their fortunes.

In addition, a wide range of nectar and / or pollen-feeding insects favour blackthorn and hawthorn blossom, notably many spring mining bees and hoverflies – plus their parasites and predators. Indeed, blackthorn blossom is a key nectar source for early spring insects, whilst hawthorn blossom attracts a wide range of diurnal insects in late spring, including a range of saproxylic beetles, and moths at night. *Aporia crataegi* itself can be expected to increase landscape-scale connectivity as pollinators (Tables 2, 7), and hence help redress concerns over the loss of pollination services in Britain (Biesmeijer *et al*., 2006).

A diverse range of invertebrates associated with *A. crataeg*i’s food plants and the structural elements of its habitats will benefit from this project, as will a number of red-listed bird species. Thus, although the focus of a species translocation project fo*r A. crataegi* would initially be on the one butterfly species, the associated landscape-scale management measures would generate a wide range of additional beneficiaries.

### *Sourcing* Aporia crataegi *stock*

The most appropriate sources of biological material of *A. crataegi* would appear to be northern France and the Pyrenees. The butterflies from these locations have the same haplotypes (evolutionary lineage) as historical specimens of the extinct British material (Todisco *et al*., 2020), they live in similar climatic conditions to those in southern-central England (Figure 9), and egg laying and larval feeding is almost exclusively confined to *Crataegus monogyna* and *Prunus spinosa* (Table 3). Distributional records from GBIF indicate clusters of records / populations in parts of Brittany and the eastern Pyrenees (yellow spots in Figure 10), where populations are reported to be large (Constanti Stefanescu, *personal communication*) and hence could provide suitable stock for translocation. Given that local adaptations may vary, establishing material from both sources over a period of two plus years would increase the genetic variation present in the newly-established population, and increase their capacity to develop adaptations suitable to British conditions.

### Assessing possible risks

Three areas that require consideration stem from the interactions of *A. crataegi* with other species. The first is the range of plant species used as larval host plants, which include a range of species in the Rosaceae, and thus includes *Malus* (apples), *Pyrus* (pears), and *Prunus* (plums, cherries). Historically, the butterfly was noted as a potential orchard pest, but it has proven difficult to track down records that distinguish between *A. crataegi* larvae as sometimes being observed on fruit trees, and the larvae having commercially significant consequences. In Europe westwards of longitude 20°E (i.e., the area we considered when reviewing the literature, and from which *A. crataegi* stock might be sourced), we found no evidence of the species being regarded as an orchard pest in the last 75 years in either the ecological or horticultural literatures. The horticultural reports we obtained for Europe from recent decades did not list *A. crataegi* as a herbivore of apples, pears, plums or cherries, not even in organic orchards. These lines of evidence (Table 3) indicate that any current use (if any) of orchards is likely to be at low levels, and effectively removed by normal horticultural practices and orchard management. Whatever the reason (genetic or horticultural practices), we conclude that *A. crataegi* is not a pest of these crops in western Europe. In contrast, it is widely reported as associated with *Crataegus* (normally *C. monogyna*) and wild *Prunus spinosa* and also with wild *Pyrus* (*P. amygdaliformis* reported) in south-eastern France and the Italian Peninsula (Table 3). The Italian populations, which were noted from cultivated fruit trees by Martelli (1931), differ in their genetics from the extinct British specimens, and from the population sources proposed for reintroduction to Britain (Todisco *et al*. 2020).

A second issue is uncertainty about the best habitat conditions and management for reestablishments to succeed. Hence, the proposed release sites should be monitored to identify rates of dispersal (emigration) from release sites, which will likely depend on the availability of shelter, nectar resources and host plants for oviposition. They should also be monitored for subsequent larval success, including their survival in the context of host plant growth forms, shelter and presence and abundance of browsing animals. Browsing may act as a source of larval mortality, especially overwinter, although this seems unlikely to prevent reestablishment, given that the New Forest, with its mix of deer, cattle and horse browsing, was a historical stronghold for the butterfly. These data will come together to identify the rates of population growth and population spread associated with different release conditions, informing subsequent releases and management options.

A third consideration is whether reintroduced butterfly stock could accidentally represent a source of new parasitoids or pathogens that could be introduced to Britain at the same time as the butterfly itself. However, of the parasitoids reported in Europe westwards of longitude 20°E (Table 1), some are generalist parasitoids across the Lepidoptera as a whole, while others are specialised primarily on *Pieris brassicae* and *Pieris rapae*, the large and small white butterfly species, respectively. Since these butterflies and their parasitoids are widespread and/or abundant in Britain already (above), it is unlikely that the reestablishment of *A. crataegi* could introduce new parasitoid species or that their establishment would alter parasitoid communities significantly. *Aporia crataegi* belongs to the butterfly family Pieridae (whites and yellows, including *P. brassicae* and *P. rapae*) and these related species are the most likely to share parasitoids and diseases. However, the risk of new ‘natural enemies’ being introduced appears slight since all of the Pieridae native to Britain already co-occur with *A. crataegi* in Europe (including in the proposed regions where stock may be sourced), and did so in Britain prior to the extinction of *A. crataegi*. Furthermore, the pierids *P. brassicae*, *P. rapae* and *Colias croceus* migrate into Britain from continental Europe every year, such that any adult-transmitted pathogen infecting the Pieridae that could establish in Britain is likely to be present already. The transport of both *Crataegus* and *Prunus spinosa* from continental Europe to Britain in recent decades also implies that any plant-associated pathogens would already have been introduced to Britain, and hence that reestablishment of *A. crataegi* would not generate any additional risks. Nonetheless, steps to ensure that introduced stock are free of parasitoids, and otherwise healthy, would be adopted. Such stock would provide opportunities for rapid population growth in the first few years (= generations) after release, before they accumulate parasitoids and pathogens shared with other species (especially from other Pieridae).

### Facilitating colonisation

Although there is potential for *A. crataegi* to traverse the English Channel occasionally, the distance is much larger than normally expected to be flown by *A. crataegi*, and this is extremely unlikely for females, which are far less mobile than males (Table 7). The chance that ‘an exceptional’ female does arrive, and does so in a location in England suitable for establishment is remote. Her offspring then need to avoid inbreeding (which can be high in relatively mobile butterfly species), which would require additional migrants to arrive at the same location in Britain within a generation or two of the first colonist. This is unlikely. The chances of establishment by spontaneous dispersal is further reduced by the lack of substantial source populations in locations where the English Channel is at its narrowest. Populations along the French coast of the English Channel are thinly distributed, and the butterfly is regarded as extinct as a breeding species in Flanders, as well as having become extinct from The Netherlands. This is believed to be associated with the intensive land use in these regions, rather than current climatic conditions (Figure 6). The only likely means by which re-establishment can be secured is by deliberate translocation (assisted colonisation).

Given the considerations reported here, it is crucial that any assisted colonisation project uses suitable genetic material associated with *Crataegus monogyna* and *Prunus spinosa*, and sourced from regions with a good climatic ‘match’. Because of the dispersal of the species (and the value in maintaining genetic diversity), hundreds to thousands of individuals need to be released across a network of sites in a well-connected landscape. Hence, the source material needs to be from large as well as suitable populations, with numbers ‘bulked up’ in captivity for a generation or two, if required.

We are in favour of monitored releases being undertaken within the landscape highlighted here to evaluate population growth, host plant use, and rates of colonisation away from release sites, aiming to develop knowledge to inform future best practices for releases. Reviewing the information presented here, we consider that there is a realistic prospect of reestablishing *Aporia crataegi* in Britain, a century after its regional extinction. As a conspicuous species able to visit flowers in gardens as well as in wilder habitats, we believe that this is likely to be widely supported by conservationists and by the public.

## ACKNOWLEDGEMENTS

This research was supported by a Research Centre Grant from the Leverhulme Trust to the Leverhulme Centre for Anthropocene Biodiversity, by the Knepp Estate and by the Knepp Wildland Foundation.

## SUPPLEMENTARY FIGURES AND TABLES

**Table S1.** Candidate sites considered for analyses of the climate match between possible reestablishment locations in England and climatic conditions across Europe.

| ID | Site | OS GridRef | Easting | Northing | Longitude | Latitude |
| --- | --- | --- | --- | --- | --- | --- |
| 1 | West Sussex | TQ136209 | 513600 | 120900 | -0.38326538 | 50.976317 |
| 2 | West Sussex | TQ144205 | 514400 | 120500 | -0.37200088 | 50.972563 |
| 3 | West Sussex | TQ146161 | 514600 | 116100 | -0.37053724 | 50.932974 |
| 4 | West Sussex | TQ165112 | 516500 | 111200 | -0.34507367 | 50.888549 |
| 5 | West Sussex | TQ168104 | 516800 | 110400 | -0.34106586 | 50.881298 |
| 6 | West Sussex | TQ095121 | 509500 | 112100 | -0.44428369 | 50.898007 |
| 7 | Devon | SX817582 | 281700 | 58200 | -3.6662995 | 50.411723 |
| 8 | Devon | SX830580 | 283000 | 58000 | -3.6479491 | 50.410186 |
| 9 | Devon | SX878546 | 287800 | 54600 | -3.5794081 | 50.380558 |
| 10 | Devon | SX874542 | 287400 | 54200 | -3.5849118 | 50.376885 |

**Figure S1.**
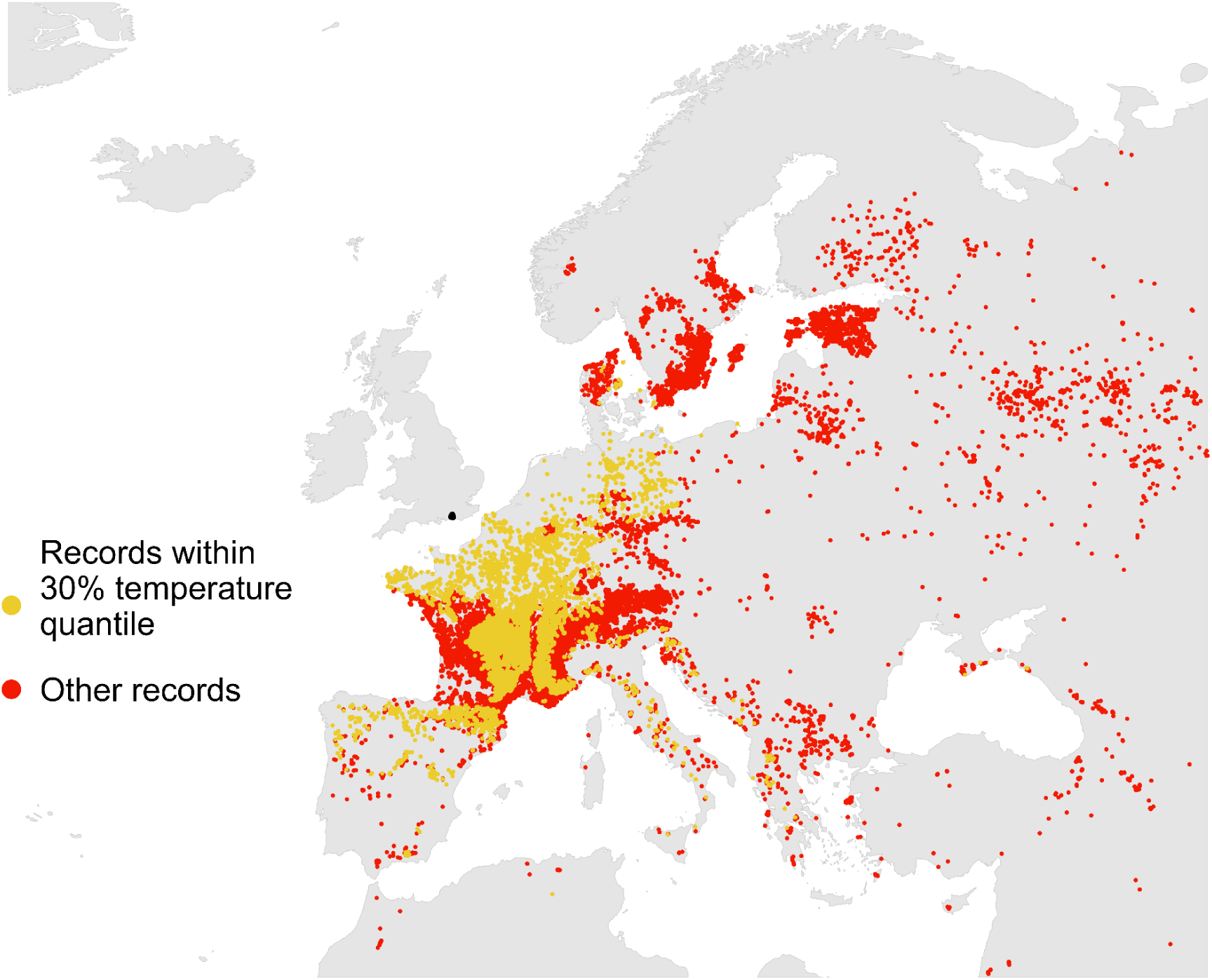
Climatic match of the West Sussex (black spot) potential reintroduction sites, illustrating areas of overlap of distributional records of the species in 2014-2023 (from GBIF) with the temperature match (30%) of the area from Figure 9b. Species records with similar (yellow) and different (red) climatic conditions are shown.

**Figure S2.**
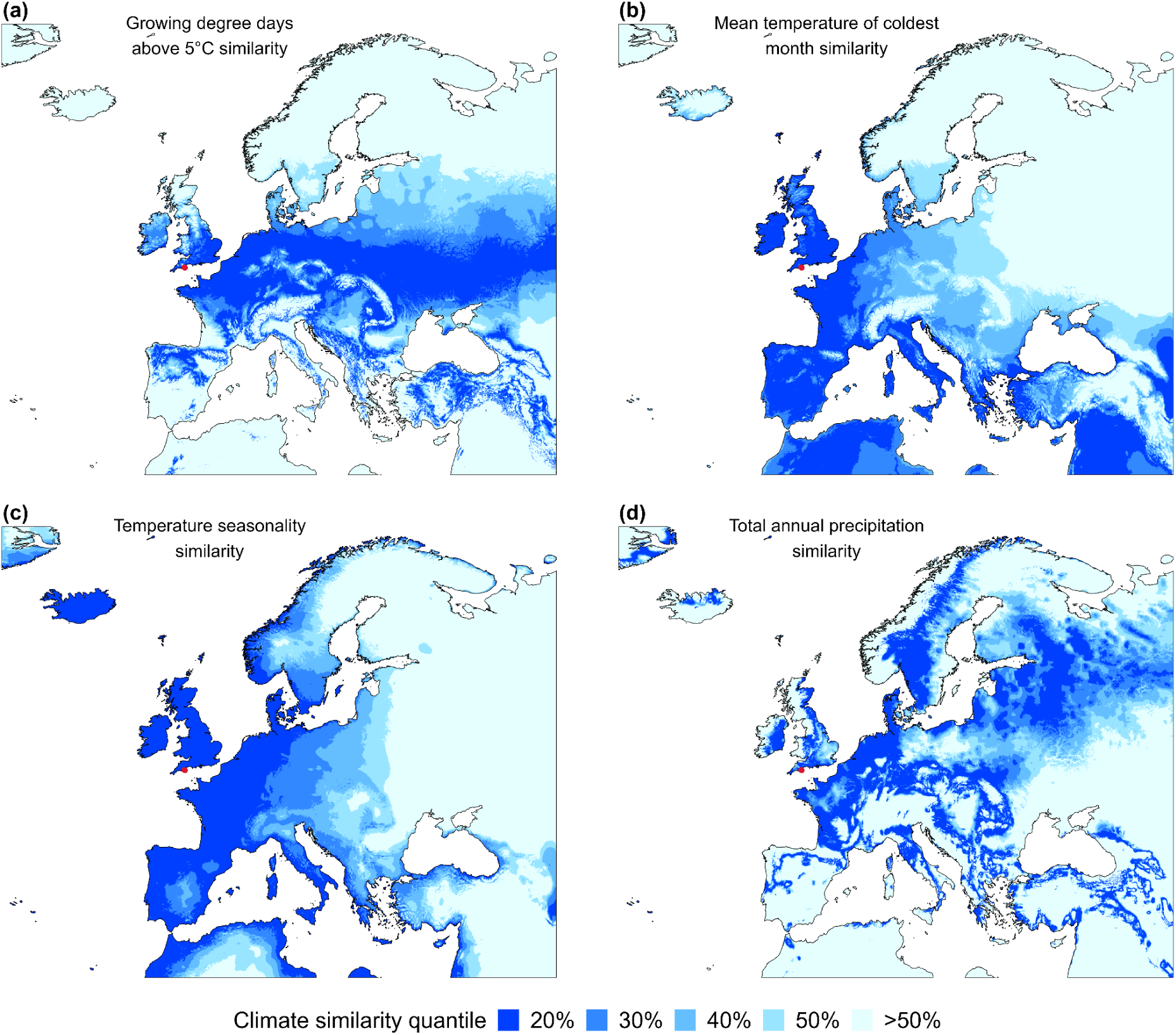
Similarity in quantiles of four climate variables to the Devon translocation sites (red dot), England.

**Figure S3.**
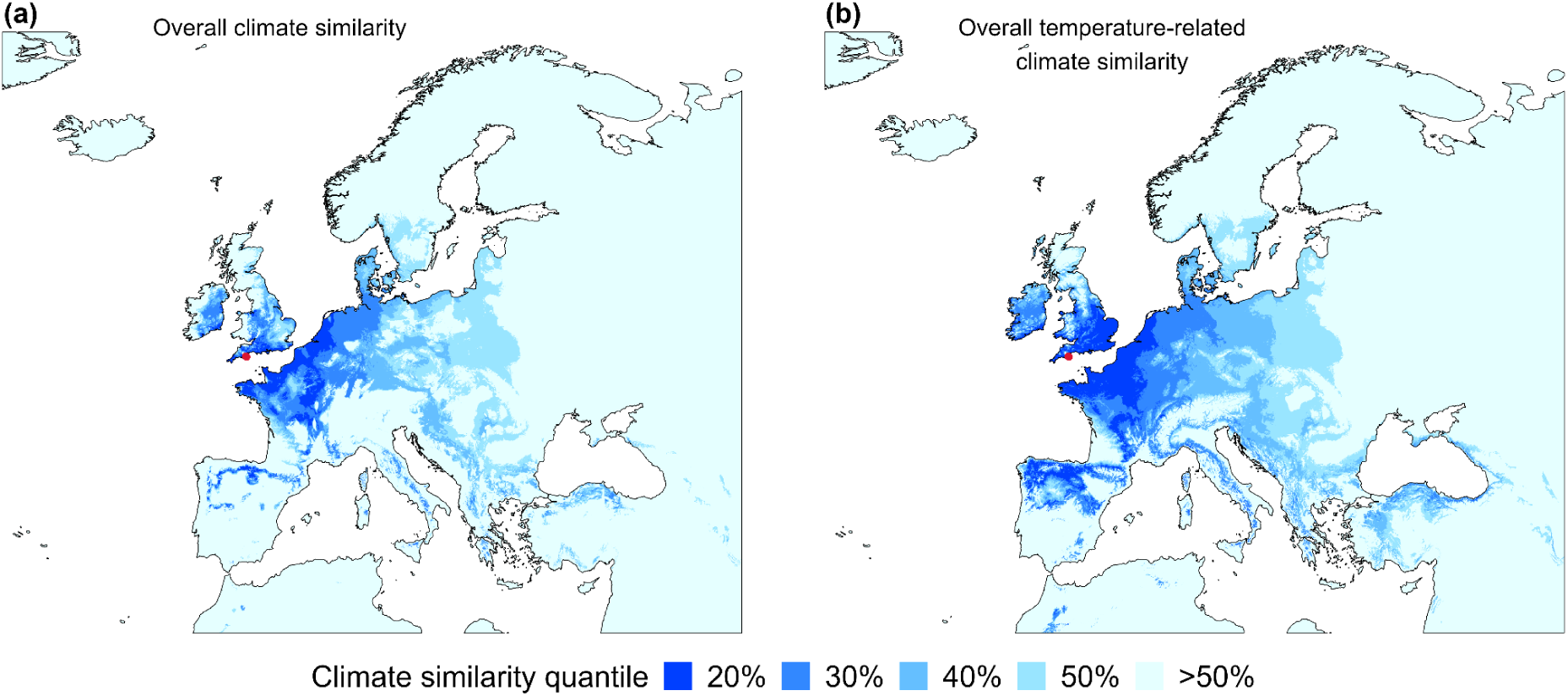
Overlap between the climatic match for all four climate variables considered (a), and for the three temperature-related variables (b), to the potential Devon translocation sites. 20% corresponds to locations in the top 20% similarity for all variables considered.

**Figure S4.**
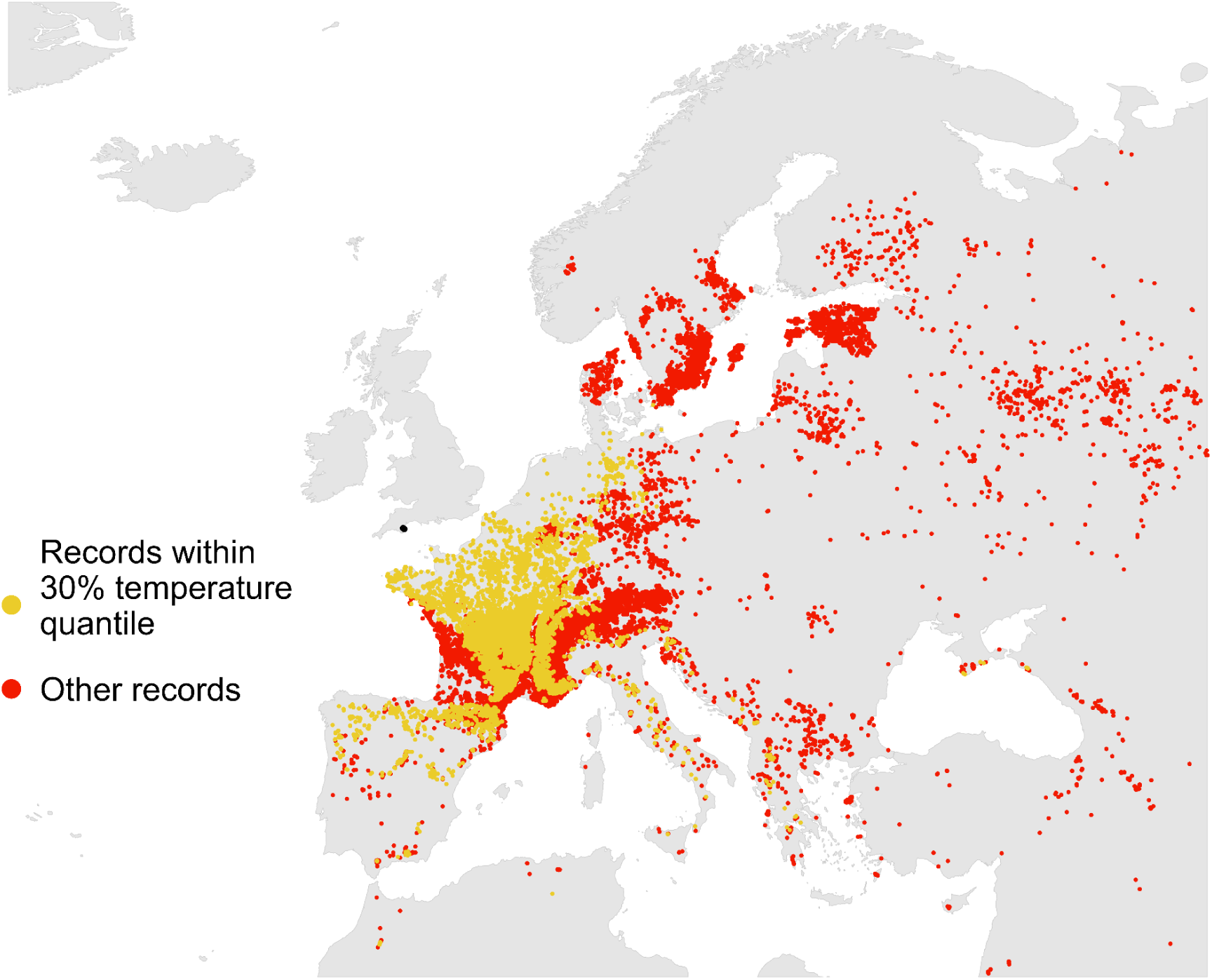
Climatic match of the Devon (black spot) potential reintroduction sites, illustrating areas of overlap of distributional records of the species in 2014-2023 (from GBIF) with the temperature match (30%) of the area from Figure S3b. Species records with similar (yellow) and different (red) climatic conditions are shown.

**Figure S5.**
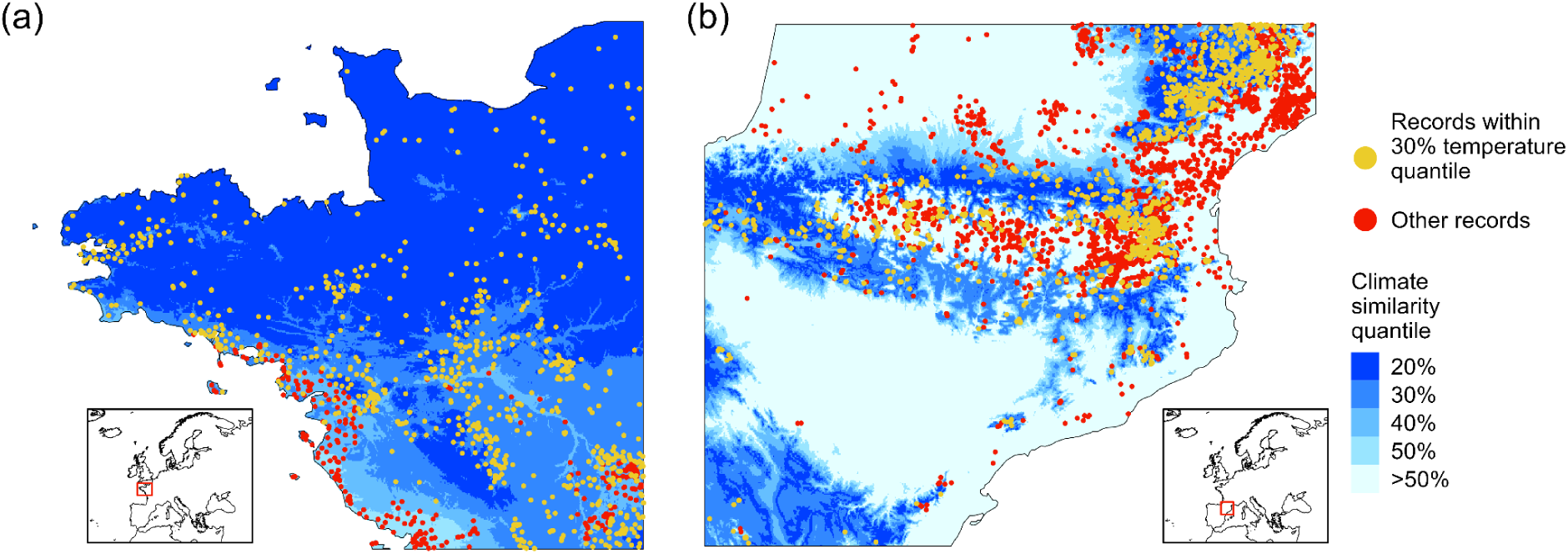
North-western France (a) and the Pyrenees (b), illustrating areas of overlap of distributional records of the species over the last decade (from GBIF) with the temperature match (30%) of the area from Figure S3b. Climate match to Devon sites. Species records with similar (yellow) and different (red) climatic conditions are shown.

## REFERENCES

Allan, P.B.M. (1948) Moths and Memories. Watkins & Doncaster,

Baker, R.R. (1969) The evolution of the migratory habit in butterflies. Journal of Animal Ecology 38: 703–746.

Baguette, M., Petit, S. and Quéva, F. (2000) Population spatial structure and migration of three butterfly species within the same habitat network: consequences for conservation. Journal of Applied Ecology 37: 100–108.

Biesmeijer, J.C., Roberts, S.P., Reemer, M., Ohlemuller, R., Edwards, M., Peeters, T., Schaffers, A.P., Potts, S.G., Kleukers, R.J.M.C., Thomas, C.D. and Settele, J. (2006) Parallel declines in pollinators and insect-pollinated plants in Britain and the Netherlands. Science 313: 351–354.

Blunck, H. and Wilbert, H. (1962) *A. crataegi* and its population changes. Zeitschrift fur Angewandte Entomologie 50: 166–221.

Broad, G., Shaw, M. and Godfray, H. (2016) Checklist of British and Irish Hymenoptera - Braconidae. Biodiversity Data Journal 4: e8151.

Caro-Miralles, E. and Gutiérrez, D. (2023) Butterfly population trends track vegetation encroachment but not climate warming in a Mediterranean mountain. Biodiversity and Conservation 32: 2017–2035.

Carroll, M.J., Anderson, B.J., Brereton, T.M., Knight, S.J., Kudrna, O. and Thomas, C.D. (2009) Climate change and translocations: the potential to re-establish two regionally-extinct butterfly species in Britain. Biological Conservation 142: 2114–2121.

Chowdhury, S., Fuller, R.A., Dingle, H., Chapman, J.W. and Zalucki, M.P. (2021) Migration in butterflies: a global overview. Biological Reviews 96: 1462–1483.

Cross, J.V., Solomon, M.G., Babandreier, D., Blommers, L., Easterbrook, M.A., Jay, C.N., Jenser, G., Jolly, R.L., Kuhlmann, U., Lilley, R. and Olivella, E. (1999a) Biocontrol of pests of apples and pears in northern and central Europe: 2. Parasitoids. Biocontrol Science and Technology 9: 277–314.

Cross, J.V., Solomon, M.G., Chandler, D., Jarrett, P., Richardson, P.N., Winstanley, D., Bathon, H., Huber, J., Keller, B., Langenbruch, G.A. and Zimmermann, G. (1999b) Biocontrol of pests of apples and pears in northern and central Europe: 1. Microbial agents and nematodes. Biocontrol Science and Technology 9:125–149.

Eeles, P, (2023) British & Irish Butterfly Rarities - Migrants, Extinctions & Introductions. Pisces Publications.

EFSA Panel on Plant Health (PLH), Bragard, C., Baptista, P., Chatzivassiliou, E., Di Serio, F., Gonthier, P., Jaques Miret, J.A., Justesen, A.F., MacLeod, A., Magnusson, C.S. and Milonas, P. (2023) Commodity risk assessment of *Crataegus monogyna* plants from the UK. EFSA Journal 21: e08003.

Fox, R., Dennis, E.B., Brown, A.F. and Curson, J. (2022) A revised Red List of British butterflies. Insect Conservation and Diversity 15: 485–495.

Fox, R., Warren, M.S. and Brereton, T.M. (2010) A new Red List of British Butterflies, Species Status No. 12, JNCC, Peterborough.

GBIF.org (23 April 2024) GBIF Occurrence Download 10.15468/dl.ek6pwj

Geervliet, J.B.F., Vet, L.E.M. and Dicke, M. (1996) Innate responses of the parasitoids *Cotesia glomerata* and *C. rubecula* (Hymenoptera: Braconidae) to volatiles from different plant-herbivore complexes. Journal of Insect Behavior 9: 525–538.

Gilbert, N. and Raworth, D.A. (2005) Movement and migration patterns in *Pieris rapae* (Pieridae). Journal of the Lepidopterists’ Society 59: 10–18.

Goded, M., Ursul, G., Baz, A. and Wilson, R.J. (2024) Changes to butterfly phenology versus elevation range after four decades of warming in the mountains of central Spain. Journal of Insect Conservation 1–15. 10.1007/s10841-024-00561-8

Hawkes, W.L., Doyle, T., Massy, R., Weston, S., Davies, K., Cornelius, E., Collier, C., Chapman, J., Reynolds, D. and Wotton, K. (2023) The most remarkable migrants–systematic analysis of the Western European insect flyway at a Pyrenean mountain pass. bioRxiv 2023.07.17.549321; doi: 10.1101/2023.07.17.549321

Hijmans, R.J., Barbosa, M., Ghosh, A. and Mandel, A. (2023) _geodata: Download Geographic Data_. R package version 0.5–9, https://CRAN.R-project.org/package=geodata

Hoegh-Guldberg, O., Hughes, L., McIntyre, S., Lindenmayer, D.B., Parmesan, C., Possingham, H.P. and Thomas, C.D. (2008) Assisted colonization and rapid climate change. Science 321: 345–346.

Hu, G., Lim, K.S., Horvitz, N., Clark, S.J., Reynolds, D.R., Sapir, N. and Chapman, J.W. (2016) Mass seasonal bioflows of high-flying insect migrants. Science 354: 1584–1587.

IUCN/SSC (2013) Guidelines for Reintroductions and Other Conservation Translocations. Version 1.0. Gland, Switzerland: IUCN Species Survival Commission.

Jaastad, G., Røen, D., Bjotveit, E. and Mogan, S. (2004) Pest management in organic plum production in Norway. In: VIII International Symposium on Plum and Prune Genetics, Breeding and Pomology 734: 193–199..

Jancke, O. (1942) New means for the control of *Aporia crataegi*. Nachrichtenblatt des Deutschen Pflanzenschutzdienstes 22: 23–24.

Jugovic, J., Crne, M. and Luznik, M. (2017) Movement, demography and behaviour of a highly mobile species: a case study of the black-veined white, *Aporia crataegi* (Lepidoptera: Pieridae). European Journal of Entomology 114:113–122.

Jugovic, J., Grando, M. and Genov, T. (2017) Microhabitat selection of *Aporia crataegi* (Lepidoptera: Pieridae) larvae in a traditionally managed landscape. Journal of Insect Conservation 21: 307–318.

Jugovic, J. and Kržič, A. (2019) Behavior and oviposition preferences of a black-veined white, *Aporia crataegi* (Lepidoptera: Pieridae). Journal of Entomological and Acarological Research 51: 10.4081/jear.2019.8108

Kan-van Limburg Stirum, P. and Kan-van Limburg Stirum, B. (2012) Braconidae: *Aporia crataegi* parasitized by *Cotesia glomerata*. Filming VarWild, Callas (watched 18 April 2024, http://www.filming-varwild.com/a-crataegi.html).

Kan-van Limburg Stirum, P. and Kan-van Limburg Stirum, B. (2014) The Black-veined White (*Aporia crataegi* Linné, 1758). Filming VarWild, Callas (watched 18 April 2024, http://www.filming-varwild.com/a-crataegi.html).

Lack, D. and Lack, E. (1951) Migration of insects and birds through a Pyrenean pass. Journal of Animal Ecology 20: 63–67.

Laing, J.E. and Levin, D.B. (1982) A review of the biology and a bibliography of *Apanteles glomeratus* (L.) (Hymenoptera: Braconidae). Biocontrol News and Information 3: 7–23.

Lind, H., Franzén, M., Pettersson, B. and Anders Nilsson, L. (2007) Metapopulation pollination in the deceptive orchid *Anacamptis pyramidalis*. Nordic Journal of Botany 25: 176–182.

Maes, D., Vanreusel, W., Herremans, M., Vantieghem, P., Brosens, D., Gielen, K., Beck, O., Van Dyck, H., Desmet, P. and Vlinderwerkgroep Natuurpunt (2016) A database on the distribution of butterflies (Lepidoptera) in northern Belgium (Flanders and the Brussels Capital Region). ZooKeys 585: 143–156. 10.3897/zookeys.585.8019.

Martelli, G.M. (1931) A Contribution to the knowledge of *A. crataegi* and of some of its parasites and hyperparasites. Bollettino del Laboratorio di zoologia generale e agraria della R. Scuola superiore d’agricoltura in Portici 25: 171–241.

Merrill, R.M., Gutiérrez, D., Lewis, O.T., Gutiérrez, J., Díez, S.B. and Wilson, R.J. (2008) Combined effects of climate and biotic interactions on the elevational range of a phytophagous insect. Journal of Animal Ecology 77: 145–155.

Met Office (2024) UK temperature, rainfall and sunshine time series https://www.metoffice.gov.uk/research/climate/maps-and-data/uk-temperature-rainfall-and-sunshine-time-series (accessed 18 April 2024).

National Climate Information Centre (2024) Met Office Hadley Centre Central England Temperature. https://www.metoffice.gov.uk/hadobs/hadcet/data/download.html (accessed 17 April 2024).

Oates, M.R. and Warren, M.S. (1990) A Review of Butterfly Introductions in Britain. Contract report for the Joint Committee for the Conservation of British Insects (JCCBI).

Pratt, C. (1983) Modern review of the demise of *Aporia crataegi* L.: the Black-Veined White. Entomologist’s Record and Journal of Variation 95: 45–52.

Quero-García, J., Iezzoni, A., Pulawska, J. and Lang, G.A., eds. (2017) Cherries: botany, production and uses. CABI.

Ratto, F. (2008) Adult behaviour and habitat availability of *Aporia crataegi* (Lepidoptera: Pieridae) in Normandy. Unpublished MSc Dissertation, Bournemouth University.

R Core Team (2023) _R: A Language and Environment for Statistical Computing_. R Foundation for Statistical Computing, Vienna, Austria. <https://www.R-project.org/>

Ryan, C. (2023) Over one million whitethorn or hawthorn imported in 2023. *Agriland*, December 14, 2023.

Shaw, B., Nagy, C. and Fountain, M.T. (2021) Organic control strategies for use in IPM of invertebrate pests in apple and pear orchards. Insects 12: 1106.

Simon, S., Brun, L., Guinaudeau, J. & Sauphanor, B. (2011) Pesticide use in current and innovative apple orchard systems. Agronomy for Sustainable Development 31: 541–555.

Solomon, M.G., Cross, J.V., Fitzgerald, J.D., Campbell, C.A.M., Jolly, R.L., Olszak, R.W., Niemczyk, E. and Vogt, H. (2000) Biocontrol of pests of apples and pears in northern and central Europe-3. Predators. Biocontrol Science and Technology 10:.91–128.

Stanković, S.S., Žikić, V., Hric, B. and Tschorsnig, H.P. (2014) Several records of Tachinidae (Diptera) reared from their hosts in Serbia and Montenegro. Biologica Nyssana 5: 71–73.

Steinhaus, E. A. (1951) Possible use of *Bacillus thuringiensis* Berliner as an aid in the biological control of the alfalfa caterpillar. Hilgardia 20: 359·381.

Thomas, C.D. and Mallorie, H.C. (1985) Oviposition records and larval foodplants of butterflies in the Atlas Mountains of Morocco. Journal of Research on the Lepidoptera 24: 76–79.

Thomas, C.D. (2011) Translocation of species, climate change, and the end of trying to recreate past ecological communities. Trends in Ecology & Evolution 26: 216–221.

Todisco, V., Vodă, R., Prosser, S.W. and Nazari, V. (2020) Next generation sequencing-aided comprehensive geographic coverage sheds light on the status of rare and extinct populations of *Aporia* butterflies (Lepidoptera: Pieridae). Scientific Reports 10: 13970.

Tresnik, S. and Parente, S. (2007) State of the art of Integrated Crop Management & organic systems in Europe,with particular reference to pest management Apple production. PAN-Europe Network, https://www.pan-europe.info/old/Resources/Reports/Apple_production_review.pdf

Tschorsnig, H-P. (2017) Preliminary host catalogue of Palaearctic Tachinidae (Diptera). Available from: http://www.nadsdiptera.org/Tach/WorldTachs/CatPalHosts/Home.html

Ubach, A., Páramo, F., Prohom, M. and Stefanescu, C. (2022) Weather and butterfly responses: a framework for understanding population dynamics in terms of species’ life-cycles and extreme climatic events. Oecologia 199: 427–439.

Van Swaay, C., Wynhoff, I., Verovnik, R., et al. (2010) IUCN Red List of Threatened Species: *Aporia crataegi*. IUCN Red List of Threatened Species. https://www.iucnredlist.org/species/159814/5339278

Verdugo Páez, A. and Verdugo Páez, M.F. (1985) Biology, ethology and distribution in the Province of Cadiz of *Aporia crataegi* Linnaeus (Lep. Pieridae). SHILAP 13: 307–311.

Whitla, R., Hens, K., Hogan, J., Martin, G., Breuker, C., Shreeve, T.G. and Arif, S. (2023) The last days of *Aporia crataegi* (L.) in Britain: evaluating genomic erosion in an extirpated butterfly. bioRxiv, 10.1101/2023.12.19.572305

Whittet, R., Cottrell, J., Cavers, S., Pecurul, M. and Ennos, R. (2016) Supplying trees in an era of environmental uncertainty: identifying challenges faced by the forest nursery sector in Great Britain. Land Use Policy 58: 415–426.

Wiklund, C. (1984) Egg-laying patterns in butterflies in relation to their phenology and the visual apparency and abundance of their host plants. Oecologia 63: 23–29.

Wilbert, H. (1959) *Apanteles glomeratus* as a parasite of *Aporia crataegi*. Beitrage zur Entomologie 9: 874–898.

Wilbert, E.L. (1960) *Apanteles pieridis* a parasite of *Aporia crataegi*. Entomophaga 5: 183–211.

Williams, C.B. (1935) Immigration of insects into the British Isles. Nature 135: 9–10.

Williams, C.B. (1951) Seasonal changes in flight direction of migrant butterflies in the British Isles. Journal of Animal Ecology 20: 180–190.

Wouters, H. (2021) Downscaled bioclimatic indicators for selected regions from 1979 to 2018 derived from reanalysis. Copernicus Climate Change Service (C3S) Climate Data Store (CDS). DOI: 10.24381/cds.fe90a594 (Accessed on 30-01-2024).

Zizka, A., Silvestro, D., Andermann, T., Azevedo, J., Duarte Ritter, C., Edler, D., Farooq, H., Herdean, A., Ariza, M., Scharn, R. and Svantesson, S. (2019) CoordinateCleaner: standardized cleaning of occurrence records from biological collection databases. Methods in Ecology and Evolution 10: 744–751.

Zografou, K., Kati, V., Grill, A., Wilson, R.J., Tzirkalli, E., Pamperis, L.N. and Halley, J.M. (2014) Signals of climate change in butterfly communities in a Mediterranean protected area. PLoS One 9: e87245.

